# ADAM17-triggered TNF signalling protects the ageing Drosophila retina from lipid droplet mediated degeneration

**DOI:** 10.1101/2020.01.09.900209

**Authors:** Sonia Muliyil, Clémence Levet, Stefan Düsterhöft, Iqbal Dulloo, Sally Cowley, Matthew Freeman

## Abstract

Animals have evolved multiple mechanisms to protect themselves from the cumulative effects of age-related cellular damage. Here we reveal an unexpected link between the TNF (tumour necrosis factor) inflammatory pathway, triggered by the metalloprotease ADAM17/TACE, and a lipid droplet (LD)-mediated mechanism of protecting retinal cells from age related degeneration. Loss of ADAM17, TNF and the TNF receptor Grindelwald in pigmented glial cells of the *Drosophila* retina leads to age related degeneration of both glia and neurons, preceded by an abnormal accumulation of glial LDs. We show that the glial LDs initially buffer the cells against damage caused by neuronally generated reactive oxygen species (ROS), but that in later life the LDs dissipate, leading to the release of toxic peroxidated lipids. Finally, we demonstrate the existence of a conserved pathway in human iPS-derived microglia-like cells, which are central players in neurodegeneration. Overall, we have discovered a pathway mediated by TNF signalling acting not as a trigger of inflammation, but as a cytoprotective factor in the retina.

## Introduction

Diseases of ageing are often caused by stress induced cellular degeneration that accumulates over time (Campisi, 2013,Lopez-Otin et al., 2013). The intrinsic causes of such damage are widespread but include toxic build-up of misfolded and aggregated proteins, as well as oxidative stress caused by the production of reactive oxygen species (ROS), which are by-products of metabolic and other physiological activity (Davalli et al., 2016,Klaips et al., 2018,Squier, 2001). Cells have evolved multiple processes to protect themselves against these potentially damaging stresses, including well characterised systems like the unfolded protein response, endoplasmic reticulum (ER)-associated degradation, and reactive oxygen species (ROS) scavenging enzymes (Bravo et al., 2013,Walter and Ron, 2011,Wellen and Thompson, 2010). Recently, lipid droplets (LDs) have begun to be implicated in the machinery of stress protection (Bailey et al., 2015,Van Den Brink et al., 2018). The significance of this role of LDs has been most studied in the fruit fly Drosophila. In central nervous system stem cell niches, elevated ROS levels induce the formation of LDs, which appear to sequester polyunsaturated acyl chains, protecting them from the oxidative chain reactions that generate toxic peroxidated species (Bailey et al., 2015). In another context, however, LDs are part of the cellular damage causing pathway: ROS generated by defective mitochondria in the Drosophila retinal neurons induces the formation of LDs in adjacent glial cells, and these LDs later contribute to glial and neuronal degeneration (Liu et al., 2015). More recent work has illustrated that toxic fatty acids produced by activated neurons in culture can be taken up by astrocytes via endocytosis, where they get subsequently catabolised by mitochondrial beta-oxidation (Ioannou et al., 2019). Although the overall significance of these mechanisms, and how they are integrated, remains to be understood, it is clear that the long-held view of LDs as primarily inert storage organelles is no longer tenable (Olzmann and Carvalho, 2019). Furthermore, beyond their role in regulating cell survival and death, LDs are increasingly found to act in other cellular processes, including acting as platforms for the assembly of viruses, and modulators of cell signalling (Boulant et al., 2008,Cheung et al., 2010,Li et al., 2014,Sandoval et al., 2014,Welte and Gould, 2017). They also form intimate contacts with ER and mitochondria (Schuldiner and Bohnert, 2017,Thiam and Dugail, 2019) and act inside the nucleus (Layerenza et al., 2013,Soltysik et al., 2019,Uzbekov and Roingeard, 2013).

In addition to the cellular mechanisms of protection against stresses, there is also protection at the level of the whole organism. This higher order, coordinated protection is primarily mediated by the inflammatory and immune systems (Chen et al., 2018,Ferrucci and Fabbri, 2018,Franceschi et al., 2018,O’Neil et al., 2018). Inflammatory signalling pathways are increasingly understood to have relevance to an extraordinary range of biology, extending far beyond classical inflammation, and including age-related damage. This is highlighted by the ever-growing list of diseases associated with inflammation including, for example, neurodegeneration, multiple sclerosis and other neurological diseases; metabolic pathologies like type 2 diabetes and obesity/metabolic syndrome; and atherosclerosis (Ferrucci and Fabbri, 2018). The signalling molecules associated with these myriad functions are equally diverse, but TNF (tumour necrosis factor) stands out as being a primary trigger of much classical and non-classical inflammatory signalling (Sedger and McDermott, 2014). Like many cytokines and growth factors, TNF is synthesised with a transmembrane (TM) anchor, and its release as an active signal is triggered by the proteolytic cleavage of the extracellular domain from its TM anchor by the ‘shedding’ protease ADAM17 (a disintegrin and metalloproteinase 17; also known as TACE, TNF alpha converting enzyme) (Black et al., 1997). Its function of being the essential trigger of all TNF signalling makes ADAM17 a highly significant enzyme. But in fact its importance is even greater because, in addition to shedding TNF, it is also responsible for the release of many other signalling molecules including EGF (epidermal growth factor) receptor ligands (Baumgart et al., 2010,Dang et al., 2013,Sahin et al., 2004). Consistent with this central role in inflammation and a wide range of other cellular events, ADAM17 has been extensively studied, both from a fundamental biological perspective and also as a drug target.

The fruit fly Drosophila, an important model organism for revealing the molecular, cellular and genetic basis of development, is increasingly used to investigate conserved aspects of human physiology and even disease mechanisms. With this motivation, we have investigated in flies the pathophysiological role of ADAM17. We report the first ADAM17 mutation in Drosophila, and find that the mutant flies exhibit abnormally high lipid droplet accumulation followed by severe age- and activity-dependent neurodegeneration in the adult retina. These observations have uncovered a new cytoprotective pathway mediated by soluble TNF acting not as a trigger of inflammation, but as a trophic survival factor for retinal glial cells. We have also shown that inhibition of ADAM17 in human iPSC-derived microglia-like cell lines leads to the same cellular phenomena as seen in fly retinas – lipid droplet accumulation, ROS production and generation of peroxidated lipids – suggesting that ADAM17 may have an evolutionarily conserved cytoprotective function.

## Results

### Loss of ADAM17 in glial cells triggers age-dependent degeneration

To investigate the full range of its pathophysiological functions, we made Drosophila null mutants of ADAM17 using CRISPR/Cas9. The knockout flies did not display any obvious defects during development and were viable as adults. They did, however, have reduced lifespan, indicating potential physiological defects. We pursued this possibility by more detailed analysis of the adult ADAM17 mutant (*ADAM17^-/-^*) retina, a tissue widely used for studying age-related degeneration of the nervous system (Morante and Desplan, 2005). The retina of the Drosophila compound eye comprises ommatidial units of eight photoreceptors, each containing a light-collecting organelle called the rhabdomere, that project axons into the brain. Photoreceptor cell bodies are surrounded by pigmented glial cells (PGCs) believed to provide metabolic support to the neurons (Edwards and Meinertzhagen, 2010,Liu et al., 2015). *ADAM17^-/-^* retinas exhibited extensive degeneration of both photoreceptor neurons and the surrounding PCGs in 5-week old flies (Figure 1A-C, L). The same result was seen with a heterozygous mutant allele in combination with a chromosome deficiency that includes the ADAM17 locus (*ADAM17^-^*/Df)(Figure 1D, L). This supports the conclusion that the degeneration was indeed caused by the loss of ADAM17. To further confirm the role of ADAM17 in the degeneration phenotype, we analysed the siRNA knockdown of ADAM17 using a retinal specific driver, *GMR-GAL4*. This too showed glial and neuronal degeneration (Figure 1E-F, L).

**Figure 1:**
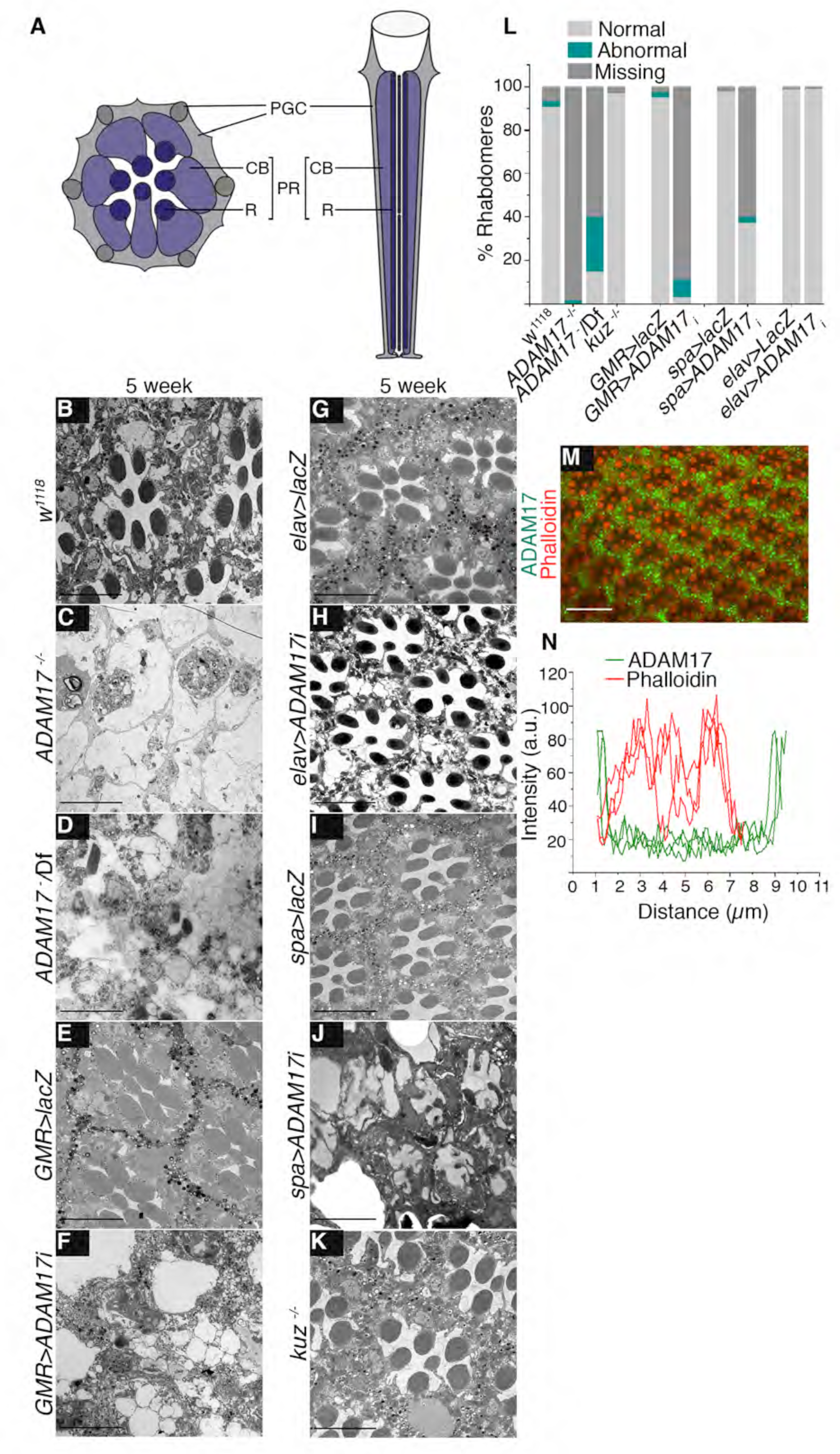
Glia specific loss of ADAM17 results in age-associated retinal degeneration. **A.** Diagram of a tangential (left) and horizontal (right) section of the adult Drosophila retina, with photoreceptors highlighted in violet and the pigmented glial cells (PGCs) in grey. PR: photoreceptor; R: rhabdomere; CB: cell body**. B-K.** Transmission electron microscopy (TEM) images of 5 week old adult retinas showing the overall structure of groups of ommatidia in (B) wildtype; (C) *ADAM17^-/-^* mutant; (D) *ADAM17^-/Deficiency^*; (E, F) RNAi against *ADAM17* and control (*lacZ*), expressed throughout the retina; (G, H) RNAi against *ADAM17* and control (*lacZ*), expressed in the neurons; (J,K) RNAi against *ADAM17* and control (*lacZ*), expressed in the pigmented glial cells; and (K) a *kuz*^-/-^ mutant. **L.** Quantitation of the percentage of normal, abnormal and degenerating rhabdomeres from TEM images for the genotypes above; n=180 ommatidia from 3 different fly retinas for each. **M.** Whole-mount retina stained with anti-ADAM17 (green) and phalloidin to mark the photoreceptor membranes (red). **N.** Line intensity profiles delineating expression of ADAM17 and phalloidin across individual ommatidia, n=10 fly retinas. Scale bar: 10μm. **See Figures S1**

An ADAM17-specific antibody revealed that expression of the enzyme is strongly enriched in the PGCs that surround the photoreceptor neurons (Figure 1M-N). This expression pattern hinted that ADAM17 function may be required specifically in glial cells, so we tested this idea by cell-type specific knock-down. ADAM17 siRNA, expressed specifically in neurons with *elav-GAL4*, produced no phenotype (Figure 1G-H, L, S1 L). In sharp contrast, glial specific knockdown with a sparkling-GAL4 driver, caused severe and characteristic degeneration (Figure 1I-J, L, S1 L). We confirmed that ADAM17 is not needed in neurons with another neuron-specific GAL4 driver, Rh1-GAL4, which drives expression specifically in photoreceptors (Figure S1A-B, K-L). The specific requirement of ADAM17 in glia was further demonstrated by showing that glial expression of ADAM17 was sufficient to rescue the degeneration phenotype of the mutant (Figure S1C-D, K). We also investigated the specificity of the phenotype by examining fly mutants of the closely related ADAM metalloprotease, Kuzbanian, the Drosophila orthologue of ADAM10 (Qi et al., 1999). *kuz^-/-^* retinas showed no signs of degeneration (Figure 1K-L).

The degeneration we observed was progressive with age. Ommatidia in the retinas of one day old *ADAM17* mutant flies were mostly intact (Figure 2A-B), although they did display enlarged PGCs, indicating some abnormalities. Retinas of two, three and four week old flies showed progressively more severe phenotypes, with only few identifiable neurons or glia seen in *ADAM17* mutants from about three weeks of age (Figure S1E-K). Collectively, these observations imply that ADAM17 in the glial cells of the adult retina protects both neurons and glia from age-dependent degeneration.

**Figure 2:**
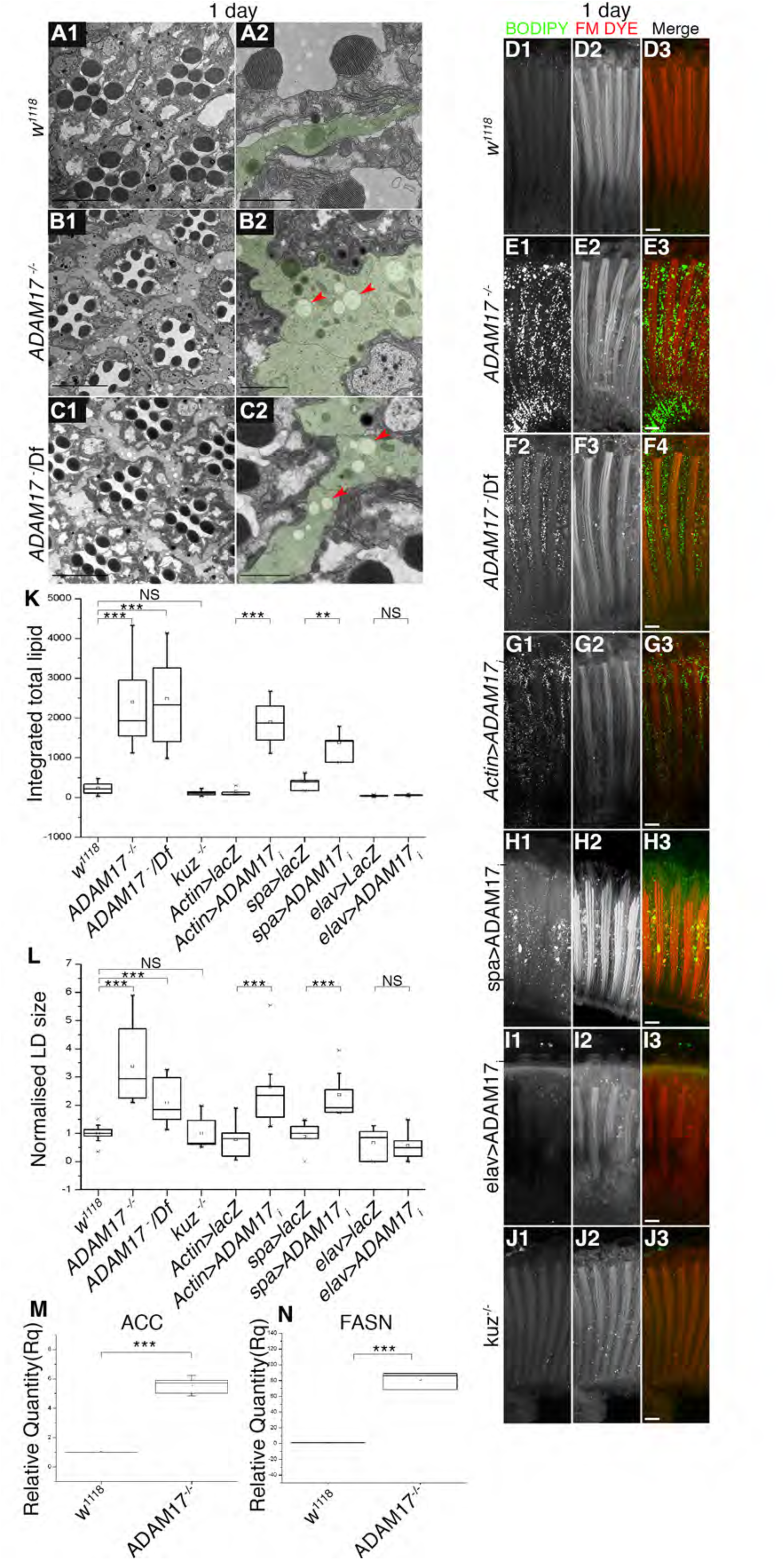
Abnormal LD accumulation in young adult retinas upon a loss of ADAM17 in glial cells. **A-C.** TEM images of 1day old adult retinas showing the overall structure of clusters of ommatidia (A1, B1, C1), or a single ommatidium (A2, B2, C2) in wild type (A1, A2), *ADAM17*^-/-^ mutant (B1, B2) or *ADAM17*^-^/Df (C1, C2). Red arrows show lipid droplets in PGCs, green shading highlights PGCs. **D-J.** Fluorescent images of 1 day old fly retinas stained with BODIPY (green) and FM-dye (red) to mark lipid droplets and the photoreceptor membranes respectively; (D) wildtype; (E) *ADAM17*^-/-^ mutant; (F) *ADAM17*^-^/Df; (G) knockdown of ADAM17 throughout the retina; (H) knockdown in glial cells; (I) knockdown in neurons; and (J) *kuz*^-/-^. n=10 fly retinas. **K-L.** Quantitation of the BODIPY signal shown as integrated total lipid and normalised LD size of lipid droplets for the genotypes in (D-I). **M-N.** Q-PCR analysis of mRNA transcripts of lipogenic genes-Acetyl CoA carboxylase (ACC) and Fatty Acid Synthase (FASN) from heads of wild type and *ADAM17*^-/-^ mutants; n=4 biological replicates. ***p<.001, **p<.01. Scale bar for A2, B2 and C2: 2μm and 10μm for all others. **See Figures S2, S3 and S4**

### Retinal degeneration of ADAM17 mutant flies is associated with LD accumulation

To investigate the fundamental cause of the retinal degeneration, we looked in more detail for the earliest detectable phenotype. We saw no visible exterior eye defects, suggesting that there was no defect in eye development (Figure S2A-B). However, ultrastructural investigation by transmission electron microscopy revealed that PGCs of one day old *ADAM17*^-/-^ and *ADAM17^-^*/Df flies were enlarged and had an accumulation of lipid droplet-like structures; these were rare in wild-type retinas (Figure 2A-C). To confirm the identity of these structures, we labelled one day old retinas with BODIPY 493/503 and FM 4-64FX dyes, which respectively label neutral lipids and cellular membranes. There was a striking accumulation of BODIPY-positive LDs in both *ADAM17*^-/-^ and *ADAM17^-^*/Df retinas (Figure 2D-F, K). In order to quantify and compare lipid accumulation between conditions, we used image analysis to measure LD number, size, and a combined measure of ‘integrated total lipid’ (see Star Methods). In comparison to wild type, we observed a clear increase in the average size of LDs in ADAM17 mutants (Figure 2L), and an even more pronounced increase in integrated total lipid (Figure 2K). LD accumulation was also caused by knockdown of ADAM17 with *Actin-GAL4*, which drives expression in both neurons and glial cells (Figure 2G, K-L). As with the degeneration phenotype, knockdown of ADAM17 specifically in PGCs, but not in neurons, also led to LD accumulation (Figure 2H-I, K-L); as before, lack of requirement in neurons was confirmed using an alternative neuronal GAL4 driver, Rh1 (data not shown). Also consistent with the degeneration phenotype, we showed that glial specific expression of ADAM17 rescued the LD phenotype of the ADAM17 mutant (Figure S3C-G); and that loss of the ADAM10 homologue, Kuzbanian, caused no accumulation of LDs in the retina (Figure 2J-L).

Increase in LD amount and size was accompanied by increased expression of lipogenic genes. We compared by qPCR the transcript levels of acetyl coA carboxylase (ACC) and fatty acid synthase 1 (FASN1), both enzymes in the lipid biosynthetic pathway. Both genes were upregulated in *ADAM17*^-/-^ fly head lysates (Figure 2 M-N). Consistent with all other data, when knocking down ADAM17 specifically in neurons or glia, we found that lipid synthesis genes only responded to glial ADAM17 loss (Figure S3H-I). Significantly, we detected no upregulation of lipogenic gene transcripts, or in the numbers of LDs, in larval brain, eye or wing imaginal discs of ADAM17^-/-^ mutants (Figure S4A-H), implying that the defects arise in adulthood, rather than being caused by earlier developmental defects. In summary, glial LD accumulation and upregulation of lipogenic genes mirrored, but preceded, cell degeneration in all experimental contexts we tested.

### Disrupting LD accumulation rescues degeneration in *ADAM17* mutants

There is a growing link between LD accumulation and the onset of neuronal degeneration (see, for example, (Liu et al., 2015,Van Den Brink et al., 2018). When we measured the temporal pattern of LD accumulation in *ADAM17^-/-^* adult retinas, we observed a sharp decline in the numbers of LDs with age, starting at around 1 week, with their almost complete disappearance by about two weeks after eclosion (Figure S4 A-H). This corresponds to the time when degeneration in the *ADAM17^-/-^* retinas was becoming prominent (Figure S1E-F, K). To explore a potential functional link between LDs and cell degeneration, we asked whether degeneration depends on prior LD accumulation. We expressed the LD associated lipase, Brummer (Bmm), a homologue of human adipocyte triglyceride lipase, either in PGCs or neurons using the same drivers as above, which are expressed from early in development. In PGCs, this continuously elevated lipase activity led to a striking reduction of LD accumulation in ADAM17^-/-^ retinas (Figure 3A-C, E-G). In contrast, lipase expressed specifically in neurons had no effect on the LD accumulation phenotype of ADAM17 loss (Figure 3D-G). Significantly, upon ageing, the flies in which LD accumulation was prevented by lipase expression showed almost complete rescue of retinal degeneration (Figure 3H-L). We conclude that prior LD accumulation in PGCs is causally linked to the degeneration induced by loss of ADAM17 in the adult retina.

**Figure 3:**
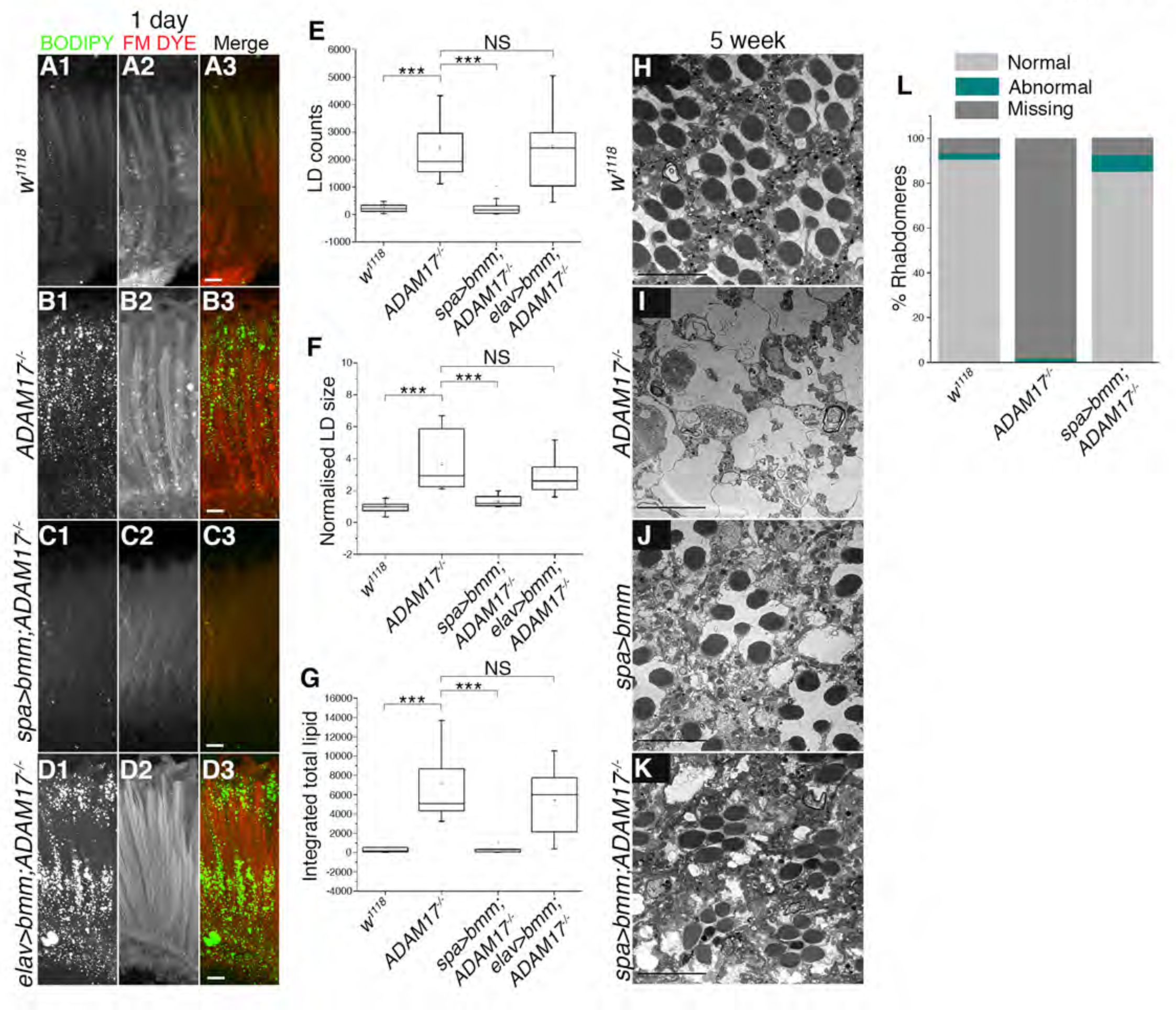
Overexpressing lipase in *ADAM17* mutant retinas rescues both lipid droplet accumulation and retinal degeneration. **A-D.** Fluorescent images of 1 day old retinas, stained with BODIPY (green) and FM-dye (red) to stain lipid droplets and the photoreceptor membranes respectively; (A) wild type; (B) *ADAM17*^-/-^ mutant; (C) overexpression of lipase in glial cells of *ADAM17*^-/-^ mutant; (D) overexpression of lipase in neurons of *ADAM17*^-/-^ mutant. **E-G.** Quantitation of the BODIPY signal shown as (E) LD numbers; (F) normalised LD size; (G) integrated total lipid, for the genotypes mentioned above, n=10 for each genotype. **H-K.** TEM images of 5 week old adult retinas corresponding to (H) wildtype; (I) *ADAM17*^-/-^ mutant; (J) overexpression of lipase in wild-type glial cells; (K) glial overexpression of lipase in the *ADAM17*^-/-^mutant. **L.** Quantitation of the percentage of normal, abnormal and degenerating rhabdomeres observed in the TEM images for the genotypes mentioned above; n=180 ommatidia from 3 different fly retinas for each. ***p<.001. Scale bar: 10μm.

### *Drosophila* ADAM17 has metalloprotease activity

Is Drosophila ADAM17 an active protease like its mammalian counterpart? It has been predicted to be, but there has been no direct demonstration of its ability to cleave substrates. Alignment of Drosophila and human ADAM17 revealed that the N-terminal pro-domain is very different between species, and that fly ADAM17 has a shorter C-terminus compared to its human counterpart (Fig. S5A). Conversely, two essential functional domains, the catalytic site and the CANDIS juxtamembrane domain, are well conserved (Figure S5A), suggesting that the Drosophila enzyme may also have metalloprotease activity. To test this, we co-expressed Drosophila ADAM17 together with alkaline phosphatase tagged Eiger, the Drosophila homologue of TNF (Igaki et al., 2002) in Drosophila S2R+ cells. Wild-type ADAM17, but not a mutant in which the protease catalytic site was defective (dADAM17HE), was able to proteolytically shed Eiger from S2R+ cells in response to stimulation by the ADAM17 activator PMA (Sahin et al., 2004). Shedding was inhibited by the broad spectrum ADAM protease inhibitor TAPI-1 (Figure 4A). We also showed that Drosophila ADAM17 cleaved a canonical mammalian ADAM17 substrate, the interleukin1 receptor IL-1R_II_ (Lorenzen et al., 2016)(Figure S4B). Since TAPI-1 could inhibit Drosophila ADAM17, we used it to test whether the ADAM17 requirement in flies was dependent on its proteolytic activity. Wild-type Drosophila reared on TAPI-1 containing food showed LD accumulation, as seen in *ADAM17^-/-^* mutants (Figure 4B-D), indicating that the proteolytic activity of ADAM17 is indeed necessary to prevent abnormal LD accumulation. These data lead us to conclude that Drosophila ADAM17 shares overlapping proteolytic activity with conventional mammalian ADAM metalloproteases, and that, as in mammals, Eiger/TNF is indeed a substrate.

**Figure 4:**
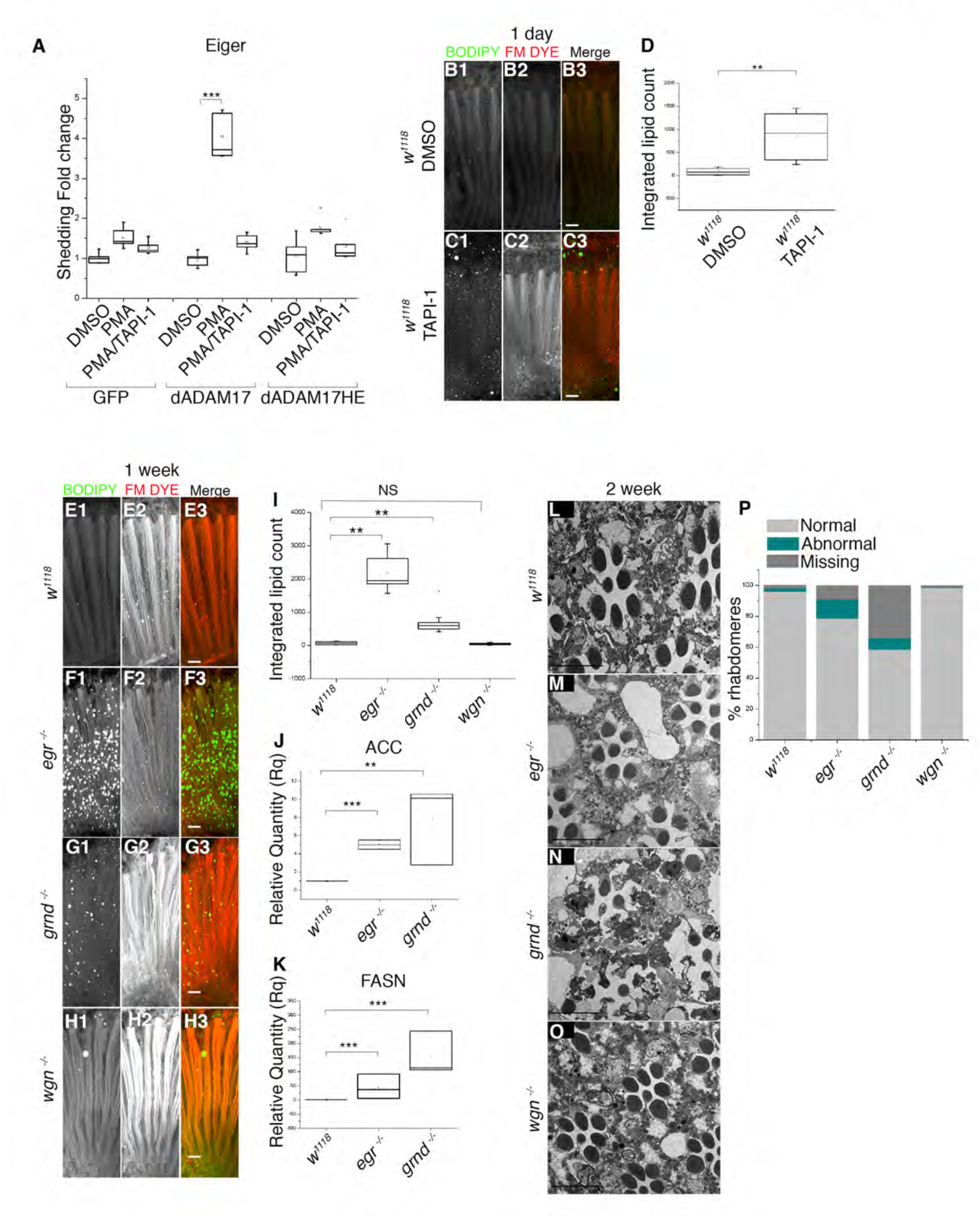
Loss of either Eiger or Grindelwald, but not Wengen, leads to abnormal LD accumulation and age-associated retinal degeneration. **A.** Alkaline phosphatase (AP)-shedding assays performed for Eiger in S2R+ cells expressing either GFP, *Drosophila* ADAM17, or activity dead mutants of *Drosophila* ADAM17, in the presence of either PMA, or PMA and TAPI-1 (DMSO is used as control; see methods); n=5. **B-C.** Fluorescent images of retinas from 1 day old flies grown on either DMSO or TAPI-1, stained with BODIPY (green) and FM-dye (red) to mark lipid droplets and photoreceptor membranes respectively. **D.** Integrated total lipid counts from retinas corresponding to flies reared either on DMSO or TAPI-1 starting from larval stages. n=10 fly retinas. **E-H.** Fluorescent images of 1 week old adult fly retinas, stained with BODIPY (green) and FM-dye (red) to mark lipid droplets and photoreceptor membranes respectively: (E) wild type; (F) *eiger*^-/-^; (G) *grindelwald^-/-^*; (H) *wengen*^-/-^. **I.** Quantitation of the BODIPY signal shown as integrated total lipid measurements for the genotypes mentioned above. **J-K.** mRNA level transcripts of ACC and FASN measured from head lysates of one week old wild type, *eiger*^-/-^ and *grindelwald*^-/-^. **L-O.** TEM images of two week old retinas corresponding to (L) control; (M) *eiger*^-/-^; (N) *grindelwald*^-/-^; (O) *wengen*^-/-^. **P.** Quantitation of the percentage of normal, abnormal and degenerating rhabdomeres from 2 week old retinas observed in TEM, corresponding to the genotypes mentioned above; n=180 ommatidia from 3 different fly retinas for each. ***p<.001, **p<.01. Scale bar: 10μm. **See Figures S5 and S6.**

### TNF pathway inhibition leads to LD accumulation and degeneration

As well as Eiger/TNF, Drosophila has two TNF receptor homologues, Grindelwald and Wengen (Andersen et al., 2015,Kanda et al., 2002,Kauppila et al., 2003). There has been some disagreement over the function of TNF signalling in flies, but among other roles, Eiger can trigger cell death (Igaki et al., 2002,Narasimamurthy et al., 2009). In another context, Eiger acts as an adipokine, released by fat body (adipocyte-like) cells and received in the brain by the Grindelwald receptor, stimulating the release of insulin (Agrawal et al., 2016). Having shown that Eiger can be shed by ADAM17, we tested whether TNF signalling mediates the function of ADAM17 in protecting retinal cells from LD accumulation and degeneration. Loss of either Eiger or Grindelwald caused an increase in LDs in the retina (Figure 4E-G, I). This was not the case with loss of Wengen, which was indistinguishable from wild type (Figure 4E, H-I). LD accumulation in *eiger* and *grindelwald* mutants was noticeably slower than in *ADAM17* mutants, visible from day 1 in about 50% of retinas but reaching full penetrance and maximum levels at about one week, instead of one day (Figure 4E-G, I). Both *eiger* and *grindelwald* mutants also showed elevated lipogenic transcripts (*ACC* and *FASN1*), although again the phenotype had a delayed onset compared to *ADAM17* mutants (Figure 4J-K). Both mutants also displayed signs of retinal degeneration by two weeks of age, and by 5 weeks full degeneration was observed (Figure 4L-N, P, S6N-Q). No retinal degeneration was seen with *wengen* mutants (Figure 4O-P).

Since Eiger is a secreted ligand and Grindelwald a receptor, they have the capacity to act in different cells. We therefore investigated in which cell types each was needed to protect cells from degeneration. Using cell-type specific RNAi, we showed that both Eiger and Grindelwald are needed specifically in glia: loss of either in PGCs, but not neurons, caused LD accumulation (Figure S6A-G), as well as the upregulation of lipogenic genes (Figure S6H-I). Knockdown of either Eiger or Grindelwald in neurons alone caused no abnormal accumulation of LDs (Figure S6G,J-L). This suggests no requirement for Eiger or Grindelwald in neurons, at least with respect to LDs. Loss of Wengen, either in glia or neurons, had no significant effect on LD accumulation or expression of lipogenic transcripts (Figure 4H, S6D-I).

Overall, these results indicate that ADAM17 protects retinal cells from LD accumulation and degeneration by shedding Eiger and subsequent activation of the TNF receptor Grindelwald, but not Wengen. Moreover, our data support a model where the ligand and the receptor are both required in glial cells.

### LD accumulation in *ADAM17* mutants depends on ROS and elevated JNK signalling

Defective mitochondria in photoreceptor neurons cause transient accumulation of LDs in neighbouring glial cells, which depends on the accumulation of reactive oxygen species (ROS) and stress signalling by the c-Jun N-terminal kinase (JNK) pathway in the photoreceptors (Liu et al., 2017,Liu et al., 2015). We therefore asked whether the retinal degeneration caused by glial ADAM17 loss also depends on ROS. We first grew larvae on food supplemented with the anti-oxidant N-acetyl cysteine amide (AD4), a ROS quencher that can cross the blood-brain barrier (Amer et al., 2008,Schimel et al., 2011). AD4 treatment rescued both LD accumulation (Figure 5A-E) and age-associated degeneration of *ADAM17^-/-^* retinas (Figure 5F-J), suggesting that ROS contribute to the *ADAM17^-/-^* phenotype. To validate this idea and identify the ROS source, we genetically quenched ROS in glia or neurons by cell-type specific expression of super oxide dismutase 2 (SOD2), a mitochondrial enzyme that destroys mitochondrially-generated ROS (Kirby et al., 2002). Quenching mitochondrial ROS specifically in glia of *ADAM17^-/-^* retinas fully rescued LD accumulation, overexpression of lipogenic transcripts, and degeneration (Figure 5K-L, O, P-R, S7D-E). Significantly lower rescue was seen with glial specific overexpression of SOD1, a cytosolic ROS quencher (Figure 5M, O, S7D-E). We also did the same experiment with catalase, another cytosolic ROS quenching enzyme. Again in contrast to mitochondrial SOD2, glial expression of catalase had little effect on LD accumulation or the induction of lipogenic genes (Figure S7A-C), supporting the conclusion that the phenotypes caused by loss of ADAM17 depends mainly on mitochondrially-generated ROS. Interestingly, neuronal specific SOD2 also caused partial rescue of LD accumulation, less complete than SOD2 expressed in glia, but significant nonetheless (Figure 5N-O). In further support for the accumulation of ROS in *ADAM17^-/-^* retinas, we performed an Oxyblot experiment to measure the extent of protein oxidation, a readout commonly used to measure cellular ROS levels (Chen et al., 2019). This showed a clear increase in *ADAM17^-/-^* mutant retinas compared to wild type (Figure 5S-T).

**Figure 5:**
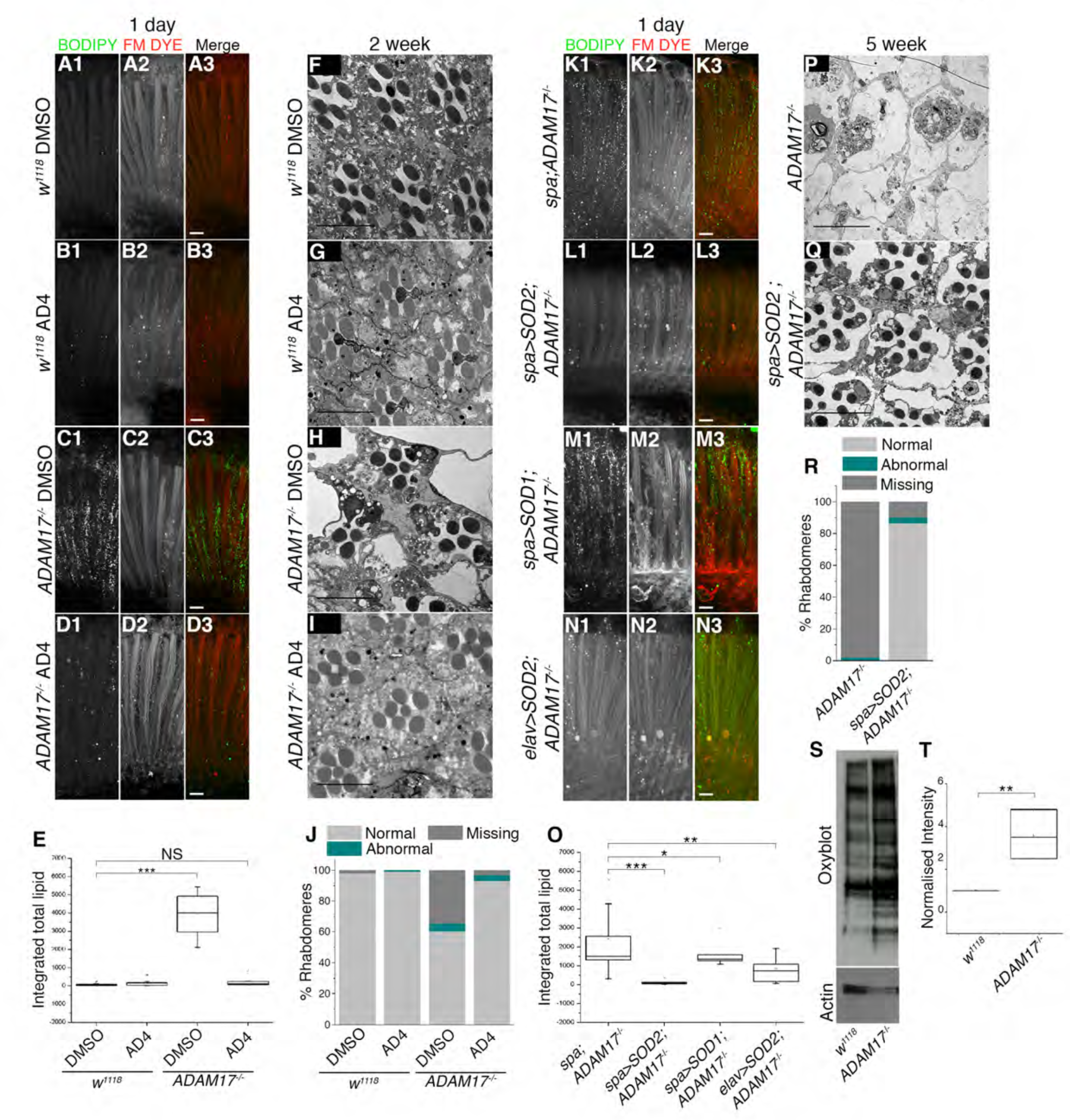
Reducing mitochondrial ROS levels in *ADAM17*^-/-^ mutants leads to a rescue of the LD accumulation and retinal degeneration. **A-D.** Fluorescent images of 1 day old adult fly retinas, stained with BODIPY (green) and FM-dye (red) to mark lipid droplets and photoreceptor membranes respectively: (A, B) wild type; (C, D) *ADAM17*^-/-^ mutants, reared either on DMSO or AD4. **E.** Integrated total lipid counts from retinas corresponding to the genotypes and treatments mentioned above; n=10 for each. **F-I.** TEM images of 2 week aged retinas corresponding to either wildtype or *ADAM17*^-/-^ mutants grown on either DMSO or AD4. **J.** Percentage of normal, abnormal or missing rhabdomeres from retinas corresponding to the genotypes and treatments mentioned above; n=3 for each. **K-N.** Fluorescent images of 1 day old adult fly retinas, stained with BODIPY (green) and FM-dye (red) to mark lipid droplets and photoreceptor membranes respectively: (K) *ADAM17*^-/-^; (L) glial specific SOD2 over-expression in an *ADAM17*^-/-^ mutant; (M) glial specific SOD1 over-expression in an *ADAM17*^-/-^ mutant; (N) neuronal specific SOD2 overexpression in *ADAM17*^-/-^ mutant. **O.** Quantitation of the BODIPY signal shown as Integrated total lipid for the genotypes mentioned above, n=10 for each. **P-Q.** TEM analysis of 5 week aged retinas corresponding to either *ADAM17*^-/-^ mutants or SOD2 over-expression in an *ADAM17*^-/-^ mutant. **R.** Percentage of normal, abnormal and missing rhabdomeres in *ADAM17*^-/-^ mutant retinas, as compared to retinas overexpressing glial specific SOD2 in an *ADAM17*^-/-^ mutant background observed with TEM; n=180 ommatidia from 3 different fly retinas for each. **S.** Oxyblot analysis of oxidative modification of proteins from retina lysates of wild type and ADAM17^-/-^. **T.** Analysis of Oxyblot intensity normalised to its respective actin controls for each genotype. Scale bar: 10μm. ***p<.001, **p<.01, *p<.05. **See Fig. S7**

Our observation that ROS quenching in neurons can partially rescue *ADAM17^-/-^* implies that, although the protective function of ADAM17 is strictly glial cell specific, and glial-generated ROS are essential to trigger LD accumulation and degeneration, neuronal ROS also contributes significantly to the phenotype. To validate and explore this involvement of neurons, we raised *ADAM17^-/-^* flies in total darkness, thereby preventing physiological ROS generation by normal photoreceptor activity. These flies exhibited almost complete rescue of degeneration and a significant, though not complete rescue of LD accumulation (Figure 6A-H). We conclude that the toxicity that leads to retinal degeneration in ADAM17^-/-^ mutants is contributed to by ROS from neighbouring photoreceptors as well as ROS generated in the glia themselves. Moreover, our data imply that the function of ADAM17 and TNF signalling in glia helps protect cells from the cumulative damage of ROS generated by normal light-induced neuronal activity.

**Figure 6:**
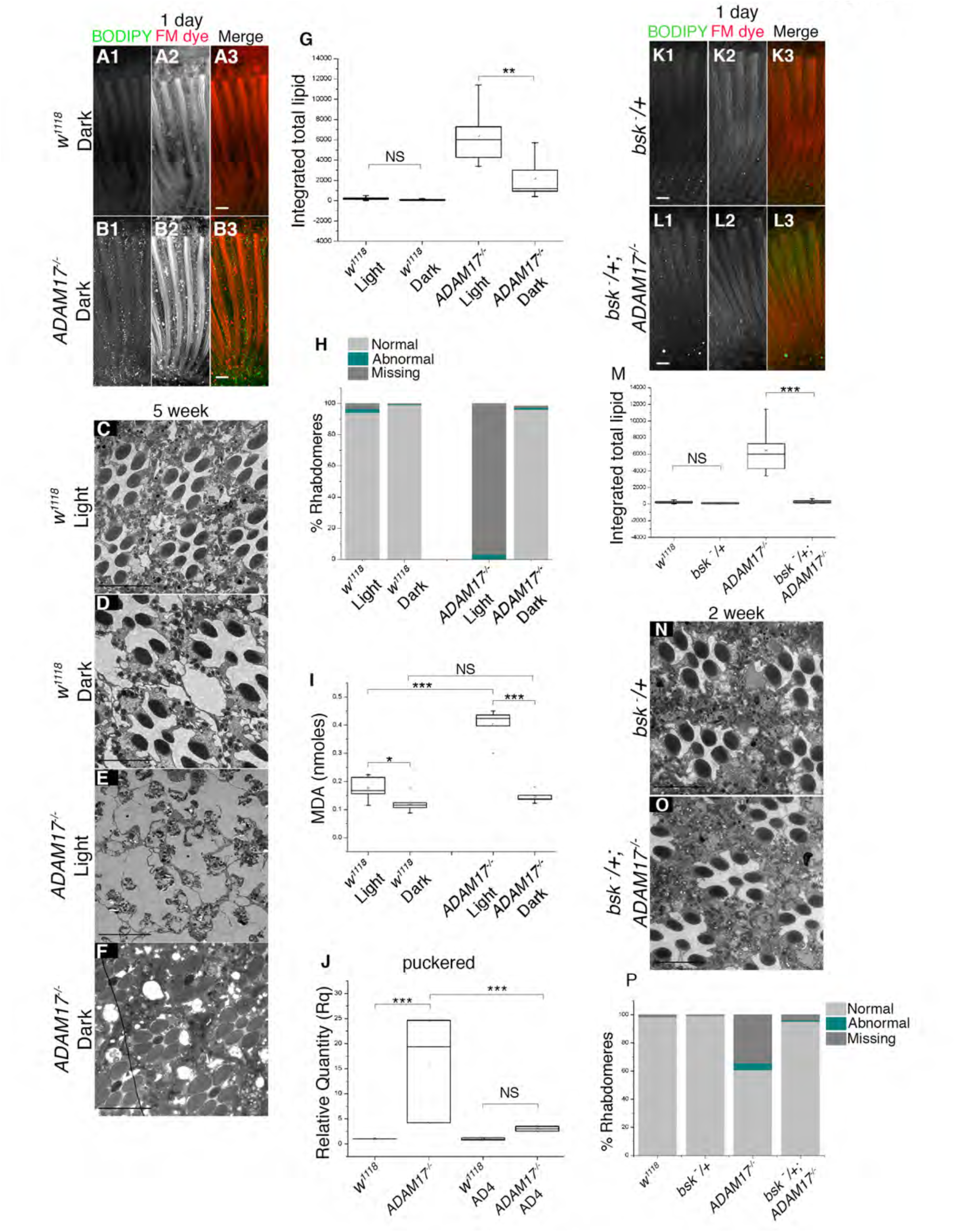
Activity dependence and role of JNK in driving LD accumulation and retinal degeneration in *ADAM17*^-/-^ mutants. **A-B.** Fluorescent images of 1 day old adult fly retinas, stained with BODIPY (green) and FM-dye (red) to mark lipid droplets and photoreceptor membranes respectively: (A) wild type; (B) *ADAM17*^-/-^, both reared in the dark. **C-F.** TEM images of 4 week old retinas: (C, D) wildtype, or (E, F) *ADAM17*^-/-^ flies reared from first larval instar either in total light (C, E) or total dark (D, F). **G.** Quantitation of the BODIPY signal shown as integrated total lipid from 1 day old retinas corresponding to the genotypes described in A-B. **H.** Percentage of normal, abnormal and missing rhabdomeres in the genotypes and treatments mentioned above. **I.** Measurement of MDA levels (in nmoles) from 1 day old retinas of wild type or *ADAM17*^-/-^ reared either in light or dark. **J.** mRNA levels of the puckered transcripts from retinas of untreated or AD4 treated wild type and *ADAM17*^-/-^ mutant retinas. **K-L.** Fluorescent images of 1 day old adult fly retinas, stained with BODIPY (green) and FM-dye (red) to mark lipid droplets and photoreceptor membranes respectively: (K) heterozygous mutant of *bsk* alone; or (L) in combination with *ADAM17^-/-^*. **M.** Quantitation of the BODIPY signal shown as integrated total lipid for the genotypes mentioned in K-L. **N-O.** TEM images of 2 week old adult retinas, showing clusters of ommatidia for a heterozygous mutant of *bsk* alone (M) or in combination with *ADAM17^-/-^* (N); n=3 for each. **P.** Percentage of normal, abnormal and missing rhabdomeres for the genotypes mentioned above; n=180 ommatidia from 3 different fly retinas for each. Scale bar: 10μm ***p<.001;, **p<.01. **See Fig. S7**

The combination of lipid accumulation and elevated ROS in *ADAM17*^-/-^ mutants suggested that toxic peroxidated lipids (Gaschler and Stockwell, 2017,Niki, 2009) might be contributing to retinal degeneration. We therefore measured the levels of malondialdehyde (MDA), a by-product and hallmark of lipid peroxidation (Chen et al., 2017). MDA was dramatically increased in *ADAM17^-/-^* retinas (Figure 6I). When rearing wild-type flies in the dark we saw a minor reduction of MDA, compared to light-reared controls (Figure 6I). This was expected and reflects the normal level of peroxidated lipids made by ROS generated by photoreceptor activity. More strikingly, the absence of neuronal activity in dark-reared *ADAM17^-/-^* flies fully rescued the accumulation of toxic peroxidated lipids (Figure 6I). This correlates well with the complete rescue of retinal degeneration seen in dark-reared flies (Figure 6E-F, H), suggesting that peroxidated lipids are indeed the major cause of cell death in *ADAM17^-/-^* mutants. This rescue of cell death by preventing photoreceptor activity supports the model that ROS contribute to the toxic effects of LDs against which ADAM17 and TNF protect glia.

We investigated whether LD formation and retinal degeneration relies on JNK signalling, a well characterised mediator of ROS-induced stress (Shen and Liu, 2006). ADAM17 loss caused a significant increase in the expression of *puckered*, a transcriptional target of the JNK pathway (Figure 6J). We also assayed phosphorylated JNK, a directed measure of JNK activity, and found it to be elevated in *ADAM17^-/-^* retinas (Figure S7F-G). The functional significance of JNK signalling was demonstrated by the observation that halving the genetic dose of JNK (the *basket* gene in Drosophila) strongly suppressed both the LD and degeneration phenotypes caused by ADAM17 loss (Figure 6K-P). This places JNK activity genetically downstream of ADAM17 loss. We investigated the relationship between ROS and JNK signalling by assaying JNK activity in retinas from flies treated with the antioxidant AD4. Quenching ROS by this means rescued elevated puckered transcript levels (Figure 6J), implying that JNK activation is triggered by ROS.

Overall, these results indicate that ADAM17 in Drosophila PGCs protects them from damage caused by ROS accumulation, which leads to the activation of the stress induced JNK pathways, LD formation, generation of peroxidated lipids, and consequent cellular degeneration.

### Loss of ADAM17 activity drives abnormal LD accumulation and mitochondrial ROS production in human iPSC-derived microglia cells

Our discovery of a new cytoprotective function for ADAM17 in Drosophila led us to question whether this role might be conserved in mammals. We therefore used human iPSC-derived microglia-like cells from three different donors, and treated them for 24 hours with the widely used metalloprotease inhibitors TAPI-1, GW280264X (GW), or GI254023X (GI). TAPI-1 is an inhibitor for ADAM17, but can also to a lesser extent inhibit other ADAMs and matrix metalloproteinases. GW inhibits specifically ADAM17 and ADAM10, while GI is a selective ADAM10 inhibitor (Chalaris et al., 2010,Moller-Hackbarth et al., 2013). Between them, they are often used to identify ADAM-dependent events and to distinguish ADAM10 from ADAM17 activity. After treatment, cells were labelled with BODIPY 493/503 and FM 4-64FX dyes to mark LDs and membrane respectively. Cells from all three donors treated with TAPI-1 or GW showed an increase in LD size and elevated total lipid, with almost no change in LD number (Figure 7A1-A3, B1-B3,C1-C3, D-E, S8A). GI treatment, specific for ADAM10, had little effect (Figure 7A4, B4, C4, D-E, S8A). These results imply that inhibition of specifically ADAM17 in these human cells causes LD defects similar to those observed in flies. We next assayed lipogenic gene expression. GW treatment of all donor cells showed an increase in the mRNA levels of FASN and LDLR, whereas GI inhibition of ADAM10 had no effect, again indicating an ADAM17-specific phenomenon (Figure 7F-G). To extend the analogy between Drosophila and human cells, we tested whether inhibiting ADAM17 activity had any effect on mitochondrial ROS levels by live imaging cells with MitoSOX. Consistent with the LD phenotype, inhibition of ADAM17, but not ADAM10, caused striking upregulation of mitochondrial ROS species (Figure 7H-K). Finally, we tested whether, as in flies, loss of ADAM17 in human cells led to an increase in the cytotoxic peroxidated lipids that we propose to be the ultimate product of the pathway. Using the dye C11-BODIPY, which undergoes a spectral shift from 590 to 520nm when bound to peroxidated lipids, we observed a sharp increase in the numbers of peroxidated LDs, only upon treatment with GW, but not with either DMSO or GI (Figure 7L-O). This result was confirmed by measuring MDA levels in untreated and treated cells. Again, a significant difference was detected in GW and TAPI-1 treated cells, with no effect upon treatment with GI (Figure S8B), demonstrating that ADAM17 inhibition causes accumulation of peroxidated lipids.

**Figure 7:**
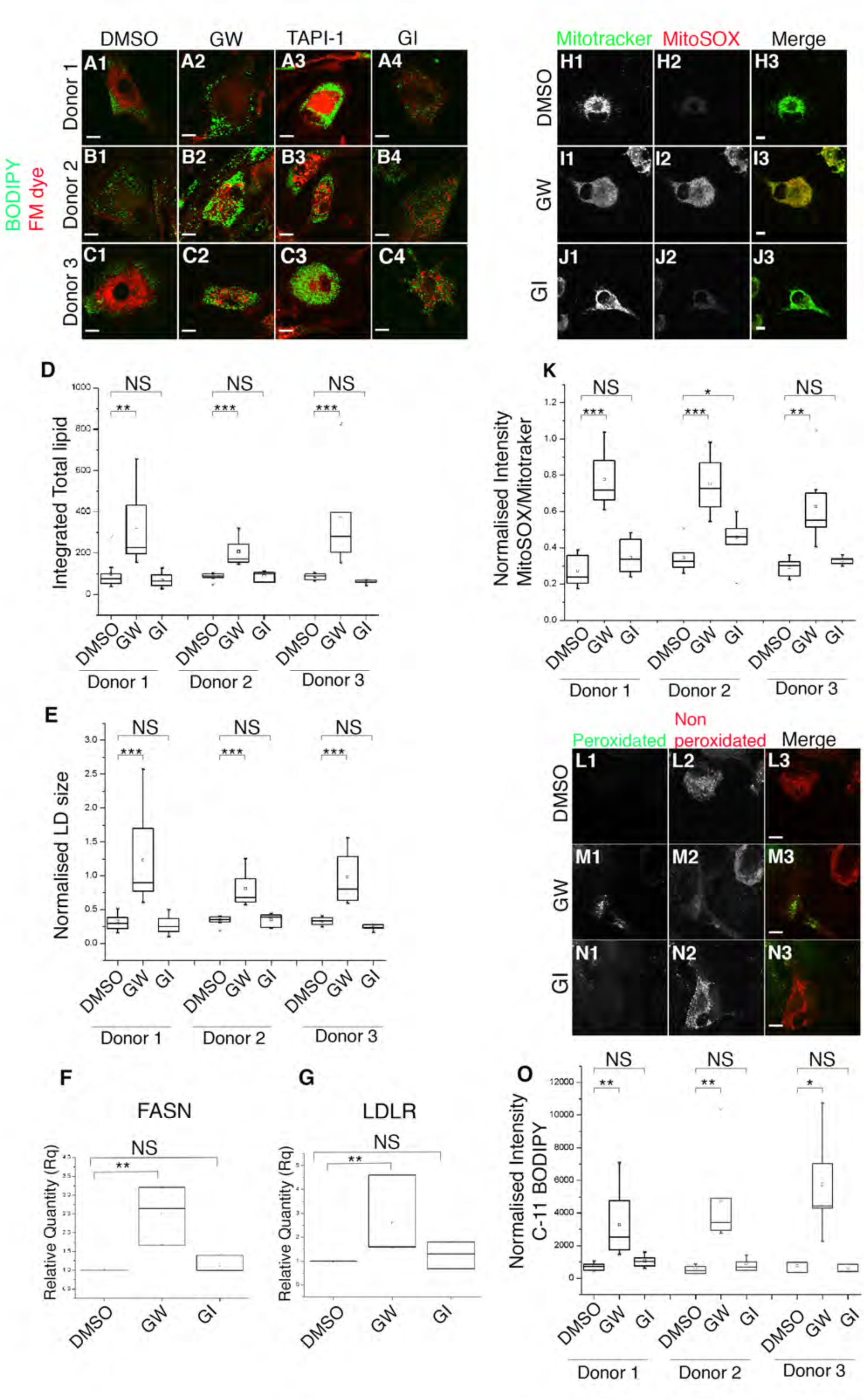
Inhibiting ADAM17 activity in human iPSC derived microglia cells leads to abnormal LD phenotypes. **A-C.** Fluorescent images of human iPSC derived microglia cells, labelled with 493/503 BODIPY (green) and FM-dye (red) to mark lipids and membranes respectively. iPSC derived microglia cells were obtained from 3 different donors, and treated with either DMSO, GW, TAPI-1 or GI. **D-E.** Quantitation of the BODIPY signal shown as integrated total lipid and normalised LD size (normalised to DMSO) for cells corresponding to each treatment and donor mentioned in A-C.. **F-G.** Q-PCR analysis of mRNA transcripts of lipogenic genes FASN and LDLR in pooled data obtained from 3 different donors of human iPSC derived microglia treated with either DMSO, GW or GI for 24hours. **H-J.** Fluorescent images of Mitotracker and MitoSOX labelling of cells from Donor no. 2 treated for 24hours with either DMSO, GW or GI. **K.** Quantitative measurements of MitoSOX intensity normalised to that of Mitotracker in iPSC derived microglia cells from three different donors, treated with either DMSO, GW or GI; n=10 cells for each treatment per donor. **L-N.** Fluorescent images of C-11 BODIPY treated microglia cells from Donor no.1, treated for 24 hours with either DMSO, GW or GI. These representative images display relative amounts of peroxidated and non-peroxidated lipids. **O.** Quantification of BODIPY C-11 staining, shown as ratios of peroxidated versus non-peroxidated lipids, depicted as normalised intensity; n=10 cells for each treatment per donor. ***p<.001, **p<.01, *p<.05. **See Fig. S8.**

In conclusion, our data show that loss of human ADAM17 leads to the same cellular phenomena in iPSC-derived human microglia as seen in Drosophila PGCs: LD accumulation, increased expression of lipogenic genes, elevated mitochondrial ROS, and abnormal levels of toxic peroxidated lipids.

## Discussion

We report here a previously unrecognised role of ADAM17 and TNF in protecting Drosophila retinal cells from age- and activity-related degeneration. Loss of ADAM17 and TNF signalling in retinal glial cells causes an abnormal accumulation of LDs in young glial cells. These LDs disperse by about two weeks after eclosion (middle age for flies), and their loss coincides with the onset of severe glial and neuronal cell death. By four weeks of age, no intact glia or neurons remain. Cell death depends on neuronal activity: retinal degeneration and, to a lesser extent, LD accumulation, is rescued in flies reared fully in the dark. We find that LD accumulation does not merely precede, but is actually responsible for subsequent degeneration, because preventing the accumulation of LDs fully rescues cell death. Our data indicate that Eiger/TNF released by ADAM17 acts specifically through the Grindelwald TNF receptor. Loss of ADAM17-mediated TNF signalling also leads to elevated production of mitochondrial ROS in glial cells, causing activation of the JNK pathway and elevated lipogenic gene expression. Together, these changes trigger cell death through the production of toxic peroxidated lipids. Importantly, toxicity is also contributed to by ROS generated by normal activity of neighbouring neurons. Finally, we show that a similar signalling module is conserved in mammalian cells: when ADAM17 is inhibited in human iPSC-derived microglial-like cells, we see the same series of events: LD accumulation, elevated mitochondrial ROS, and high levels of toxic peroxidated lipids (Figure 8).

**Figure 8:**
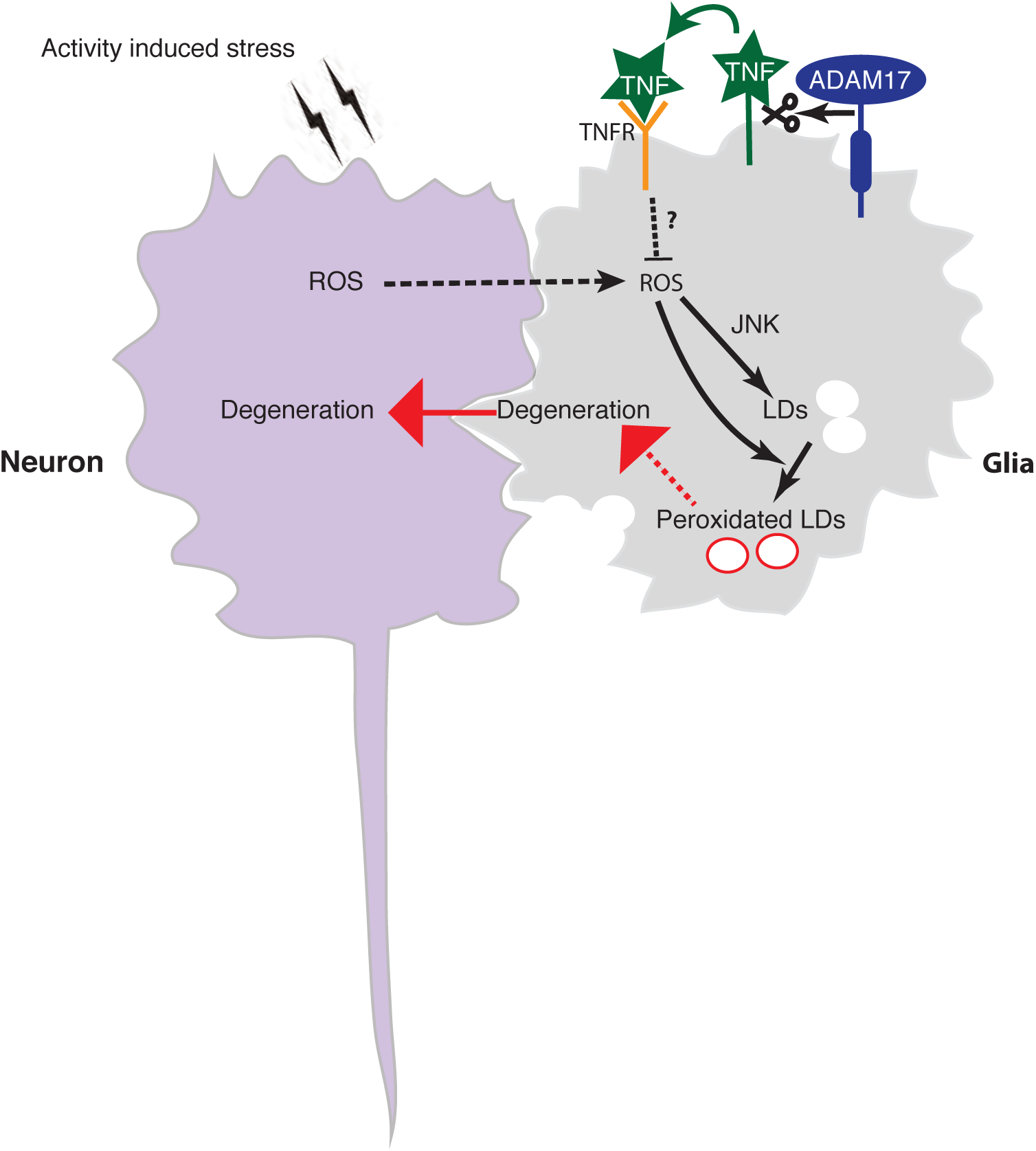
Regulation of ROS and peroxidated LDs by ADAM17 and components of the TNF pathway. Cleavage of full length TNF/Eiger by ADAM17 leads to the production of soluble TNF/Eiger, which can engage the TNF receptor, Grindelwald. TNF signalling inhibits the production of mitochondrial ROS inside glial cells, through unknown mechanisms. In the absence of TNF signalling, glial generated ROS combines with ROS from neighbouring neurons to trigger production of LDs. Upon their later dispersal, the combination of fatty acids and ROS leads to high levels of toxic peroxidated lipids, which are toxic to the glia. Finally, glial degeneration is followed by neuronal degeneration.

We propose that TNF is an autocrine trophic factor that protects retinal pigmented glial cells from age-related cumulative damage caused by the ROS that are normal by-products of neuronal activity. This ADAM17/TNF protection system is located specifically in retinal glial cells, but its role is to protect both glia and neighbouring neurons (Fig. 4G). In the absence of this TNF cytoprotective pathway, we see severe early-onset retinal neurodegeneration. Our data imply that cells die by being overwhelmed by toxic peroxidated lipids when abnormal accumulations of LDs disperse (Fig. 4G). This occurs in *Drosophila* middle age, when LDs stop accumulating and begin to disperse, triggering the cytotoxic phase of the *ADAM17*^-/-^ phenotype. It is important to emphasise that despite the ADAM17/TNF protection system being located specifically in retinal glial cells, there is neuronal involvement. Not only does TNF indirectly protect against neurodegeneration, but photoreceptor neurons are also significant sources of the ROS that generate the toxic peroxidated lipids in glia. More generally, this work provides a model for investigating more widely the functional links between ageing, cellular stress, lipid droplet accumulation and neurodegeneration. Indeed, in the light of our discovery that the pathway we have discovered in *Drosophila* is conserved in human microglia-like cells, it is significant that lipid droplets have been reported to accumulate in human microglia, cells that are increasingly prominent in the pathology of Alzheimer’s Disease and other neurodegenerative conditions (Bastos et al., 2017,Karran and De Strooper, 2016,Shinozaki et al., 2014).

In a *Drosophila* model of neuronal mitochondrionopathies, abnormal neuronal ROS production led to elevated neuronal lipid production, followed by transfer of the lipids to the PGCs, where LDs accumulated (Liu et al., 2017,Liu et al., 2015). The ADAM17 mutants we report are different, showing no evidence for abnormal lipid production in neurons. Instead, in our paradigm where TNF protects against neurodegeneration, the photoreceptor neurons are also significant sources of the ROS that generate the toxic peroxidated lipids in glia. Despite these differences between this work and what has been previously reported, a growing body of work points to a close coupling between ROS, lipid droplets and cellular degeneration (Liu et al., 2017,Liu et al., 2015,Van Den Brink et al., 2018), a relationship conserved in mammals ((Liu et al., 2015); this work).

It has become clear that LDs are much more than simply passive storage vessels for cellular lipids; they have multiple regulatory functions (Welte and Gould, 2017) Indeed, although we here highlight a developing picture of an LD/ROS-dependent trigger of cell death, in other contexts LDs have protective functions against oxidative damage, both in flies and mammals (Bailey et al., 2015,Bensaad et al., 2014).This may occur by providing an environment that shields fatty acids from peroxidation by ROS and/or by sequestering toxic peroxidated lipids. Although this superficially appears to contradict the theme of LD/ROS toxicity, it is important to recall that in LD-related cell death is not simultaneous with LD accumulation. In fact, degeneration temporally correlates with the dispersal of LDs in middle age, rather than their earlier accumulation. Together, the strands of evidence from several studies suggests that it is the combination of elevated ROS and the dispersal of abnormally high quantities of lipids from previously accumulated LDs that trigger death. This suggests that cells die by being overwhelmed by toxic peroxidated lipids when abnormal accumulations of LDs break down in the presence of high levels of ROS. Our experiments with Brummer lipase are consistent with this idea: the Brummer lipase was expressed from early in development, thereby preventing abnormal LD accumulation, and this protected against cell death. This sequence of events implies the existence of a metabolic switch, when LDs stop accumulating and begin to disperse, triggering the toxic phase of ADAM17 loss. It will be interesting in the future, and may provide insights into the normal ageing process, to understand the molecular mechanism of this age- and/or activity-dependent change.

ADAM17 is one of the most important shedding enzymes in humans, responsible for the proteolytic release of a vast array of cell surface signals, receptors and other proteins (Boutet et al., 2009,Lorenzen et al., 2016,Zunke and Rose-John, 2017). Because of its role in signalling by both TNF and ligands of the EGF receptor it has been the focus of major pharmaceutical efforts, with a view to treating inflammatory diseases and cancer (Arribas and Esselens, 2009,Mustafi et al., 2017). It is therefore surprising that it has been very little studied in Drosophila. This is the first report of Drosophila ADAM17 mutants. Here we also confirm for the first time that Drosophila ADAM17 is indeed an active metalloprotease, able to shed cell surface proteins including the Drosophila TNF homologue Eiger. The only other description of Drosophila ADAM17 function is mechanistically consistent with our data, despite relating to a different physiological context. In that case, ADAM17 was shown to cause the release of soluble TNF from the fat body so that it can act as a long range adipokine (Agrawal et al., 2016). We cannot rule out other ADAM17 substrates in different developmental or physiological contexts, although the relatively subtle phenotype of null mutant flies implies that ADAM17 does not have essential functions that lead to obvious defects when mutated. Moreover, we have not observed developmental defects or LD accumulation in any neuronal or non-neuronal ADAM17^-/-^ larval tissues, suggesting that the mechanism we report here is both age and tissue specific.

Although TNF is sometimes viewed as a specific cell death promoting signal, and the pathways by which it activates caspase-induced apoptosis have been studied extensively in flies and mammals (Aggarwal, 2003,Wajant et al., 2003) (Wallach et al., 2002), the response to TNF is in fact very diverse, depending on the biological context (Postlethwaite and Seyer, 1990,Puls et al., 1999). Indeed, its most well studied role in mammals is as the primary inflammatory cytokine, released by macrophages and other immune cells, and triggering the release of other cytokines, acting as a chemoattractant, stimulating phagocytosis, and promoting other inflammatory responses (Bradley, 2008,Sedger and McDermott, 2014). To our knowledge, however, it has not previously been shown to have trophic activity, protecting cells from stress-induced damage, in the nervous system or elsewhere.

In conclusion, our work highlights three important biological concepts. The first is to identify a new function for the ADAM17/TNF pathway in a cytoprotective role that protects *Drosophila* retinal cells against age- and activity-dependent degeneration. This contrasts with its well-established roles in inflammation and cell death. Secondly, we highlight the existence of a glia-centric cellular pathway by which the breakdown of accumulated LDs and ROS together participate in promoting stress-induced and age-related cell death. Finally, we have shown that the core phenomenon of the ADAM17 protease, acting to regulate the homeostatic relationship between ROS and LD biosynthesis, is conserved in human microglial cells, which themselves are intimately involved in neuroprotection.

## Materials and Methods

### Fly Genetics

#### Generation of *ADAM17^-/-^* flies

*ADAM17-/-* flies were generated through CRISPR using the guides <u>CTGTGGCCTCGCAATAATCT**CGG and**</u> GCC<u>TTTT</u>CGTCGAGAATTG**CGG** for targeting the first exon of the gene. These guides were microinjected into *y^1^ P(act5c-cas9, w^+^) M(3xP3-RFP.attP)ZH-2A w\** (Cambridge microinjection facility), and then subsequently screened using Melting curve analysis.

#### Transgenic flies

To generate the UAS-ADAM17 WT fly lines, the *pUASattB-ADAM17 WT* construct was introduced into the germ line by injections in the presence of the PhiC31 integrase and inserted in the attp40 landing site on the 2^nd^ chromosome (Cambridge microinjection facility).

The following lines were from the GD or KK collections of the Vienna Drosophila RNAi Centre (VDRC): *ADAM17* RNAi (v2733), *grnd* RNAi (43454), *wengen* RNAi (58994), *kuz* RNAi (v107036).

The *w^1118^*, *elav-Gal4 (8765)*, *actin-GAL4*(BL25374), *sparkling-GAL4* (BL 26656), *rh1-GAL4* (BL 68385), *gmr-GAL4* (BL9146), 54C-*GAL4* (27328), *eiger* RNAi(58993), *grnd^Minos^* ^(Mi(Baumgart et al.)^CG10176MI05292, *ADAM17 Deficiency* (27366), *UAS-bmm* (76600), *UAS-lacZ* (1776), *bsk*^1^ (3088), *UAS-SOD2* (24494), *UAS - SOD1* (24491) and *UAS-catalase* (24621) were obtained from the Bloomington Stock centre.

All crosses were performed at 29 °C in light, unless otherwise stated.

#### A detailed list of fly genotypes is presented below

##### Fig 1

*w^1118^*

*;+/+;ADAM17^-/-^*

*;+/+;ADAM17^-^/Df*

*;GMR-GAL4/+*; *UAS-lacZ/+*

*;GMR-GAL4/+*; *UAS-ADAM17_i_/+*

*;elav-GAL4/+*; *UAS-lacZ/+*

*;elav-GAL4/+*; *UAS-ADAM17_i_/+*

*spa-GAL4*;+/+;*UAS-lacZ/+*

*spa-GAL4*;+/+; *UAS-ADAM17_i_/+*

*;kuz*^-/-^

##### Fig. 2

*w^1118^*

*;+/+;ADAM17^-/-^*

*;+/+;ADAM17^-^/Df*

*;+/+*;*Actin-GAL4*/*UAS-lacZ*

*;+/+*;*Actin-GAL4*/*UAS-ADAM17_i_*

*spa-GAL4*;+/+;*UAS-lacZ/+*

*spa-GAL4*;+/+;*UAS-ADAM17_i_/+*

*;elav-GAL4/+*;*UAS-lacZ/+*

*;elav-GAL4/+*;*UAS-ADAM17_i_/+*

*;kuz*^-/-^

##### Fig. 3

*w^1118^*

*;+/+;ADAM17^-/-^*

*spa-GAL4*;UAS-bmm/+;

*spa-GAL4*;UAS-bmm/+;*ADAM17^-/-^*

*,elav-GAL4/*UAS-bmm;*ADAM17^-/-^*

##### Fig. 4

*w^1118^*

*;egr*^-/-^

*;grnd^-/-^*

*wgn^-/-^*

##### Fig. 5

*w^1118^*

*;+/+;ADAM17^-/-^*

*spa-GAL4*;+/+;*ADAM17^-/-^*

*spa-GAL4*; UAS-SOD2/+;*ADAM17^-/-^*

*spa-GAL4*; UAS-SOD1/+;*ADAM17^-/-^*

*;elav-GAL4*;UAS-SOD2/+; *ADAM17^-/-^*

##### Fig. 6

*w^1118^*

*;+/+;ADAM17^-/-^*

*;bsk^-^/+*

*;bsk^-^/+; ADAM17^-/-^*

### Electron microscopy

Samples were fixed in 2.5% Glutaraldehyde + 4% PFA + 0.1% tannic acid in 0.1M PIPES pH 7.2, for 1 hour at room temperature (RT) and then overnight at 4 °C. Samples were then washed with 0.1M PIPES at RT over the course of 2 hrs with several solution changes, including one wash with 50 mM glycine in 0.1M PIPES to quench free aldehydes. Samples were then incubated in 2% osmium tetroxide + 1.5% potassium ferrocyanide in 0.1M PIPES for 1hr at 4 °C with rotation. Post fixation, samples were washed 3 times with MQ water for 10min each. This was followed by tertiary fixation with 0.5% uranyl actetate (aq.) at 4 °C overnight. All subsequent steps were performed on a rotator set to medium-high speed. Samples were washed 3 times with MQ water and then sequentially dehydrated for 10 min each in ice cold 30%, 50%, 70%, 100% ethanol twice, anhydrous ice-cold acetone and finally in anhydrous acetone at RT. Samples were then infiltrated at RT with Durcupan resin as follows: 25% resin in acetone for 3 hrs, 50% resin in acetone overnight, 75% resin for 2-3 hrs, 100% resin for ∼6 hrs, 100% resin overnight, 100% resin 7 hrs, 100% resin overnight and 100% resin 4-5 hrs. After each change into fresh resin, the samples were spun for 30 secs in a minifuge at 6000rpm to aid infiltration. Samples were then embedded in the fresh resin in flat moulds, and cured at 60 °C for 48 hours.

Eyes were sectioned tangentially using a Leica UC7 ultramicrotome. Ultrathin (90 nm) sections were obtained using a Diatome diamond knife and transferred to formvar-coated copper or copper/palladium 2×1mm slot grids. Sections were post-stained for 5 mins with Reynold’s lead citrate and washed 3 times with warm water, blotted and air dried.

Grids were imaged on either a FEI Tecnai 12 Transmission Electron Microscope (TEM) at 120 kV using a Gatan OneView camera or, for low magnification mapping, on a Zeiss Sigma 200 Field Emission Gun Scanning Electron Microscope (FEG-SEM) at 20 kV using the STEM detector.

Fixation and dehydration for Scanning Electron Microscopy (SEM) were carried out similarly to those for the TEM. After dehydration through an ascending series of ethanols ending in 100% ethanol, samples were dried using critical point drying (CPD), without introducing surface tension artifacts. After drying, the samples were carefully mounted on an aluminum stub using silver paint. Samples were then introduced into the chamber of the sputter coater and coated with a very thin film of gold before SEM examination. All SEM samples were imaged on the Zeiss Sigma 200 Field Emission Gun Scanning Electron Microscope (FEG-SEM) at 20 kV.

### Degeneration Measurements

Missing, abnormal or missing rhabdomeres were counted manually on TEM images from a total of 60 ommatidia per retina using Image J. These counts were then averaged over three retinas per genotype. The average percentage for each genotype is displayed as a stacked column graph.

### Whole mount staining of *Drosophila* retinas

For LD staining, *Drosophila* heads were cut in half and the brain was removed to expose the retina underneath. Retinas were fixed in 3.7% formaldehyde (Sigma, F8775-500ML) in PBS for 15 min and washed 3 times in PBS1X. Retinas were incubated for 10min in BODIPY™ 558/568 (Invitrogen, D3835) to label lipid droplets, and FM™ 4-64FX (Invitrogen, F34653) to stain membranes. Retinas were then mounted in Vectashield (Vector Laboratories Ltd, H-1000) and imaged on Olympus FV1000.

### Lipid Droplet Measurements

Lipid droplet numbers and size were measured from the BODIPY signal, using an algorithm developed on Image J. Integrated total lipid was calculated as a product of normalised size and total lipid droplet numbers across an individual retina.

Normalised LD size was calculated by dividing the size of LDs across all genotypes with that of of its corresponding control.

### Immunostaining of whole mount *Drosophila* retinas

For immunostaining, retinas were dissected similarly as above. Retinas were fixed in 4% paraformaldehyde (VWR, 43368.9M) in PBS for 15 min and washed 3 times in PBS1X-triton0.3%. Retinas were labelled with anti-ADAM17 (Abcam) overnight at 4 °C followed by secondary antibody and Phalloidin staining (life technologies, A12380) for 2h at room temperature. Retinas were mounted in Vectashield (Vector Laboratories Ltd, H-1000) and imaged on Olympus FV1000.

### Line Intensity profiles

Line intensity profiles for ADAM17 and Phalloidin were calculated manually using the Plot profile plugin on Image J. Intensity values were measured along a line in FIJI averaging 10um,that spanned a single ommatidia, and plotted along its distance.

### mRNA extraction and quantitative PCR

20 frozen fly heads were used to extract RNA using Trizol reagent with the help of a Direct-zol mRNA kit, according to manufacturer’s instructions. cDNA was prepared from 0.4µg RNA using the qpcrbiosystems kit (PB30.11-10). Resulting cDNA was used for Quantitative PCR (qPCR) combining it with Taqman Gene Expression Master Mix (Applied Biosystems) and Taqman probes (all Thermo Fischer). qPCR was performed on a StepOnePlus system Thermocycler (Applied Biosystems). Each biological experiment was carried out with three independent technical replicates and normalised to Actin in each case. Error bars indicate the standard error from mean for at least three different biological replicates. Taqman probes used for all experiments are as follows: Drosophila ACC (Dm01811991_m1), Drosophila FASN (Dm01821412_m1), Drosophila TACE/ADAM17(Dm02146367_g1), Drosophila actin (Dm02362162_s1), Human FASN (Hs01005624), Human LDLR (Hs00181192_m1) and Human Actin (Hs01060665_g1).

### Statistical Analysis for LD and Q-PCR measurements

All datasets were analysed in Prism. Normality and homogeneity of variance were used to determine whether the data met the assumptions of the statistical test used. All datasets were assumed to be independent. Datasets with unequal variance were analysed using the Kruskal Wallis test followed by Dunn’s test for post hoc analysis for significance due to unequal sample sizes. All other datasets were quantified for significance using Student’s t test. Significance is defined as *p < 0.05, **p<.01, ***p<.001 and error bars are shown as standard error of the mean (SEM) unless otherwise noted. For fly experiments, more than 10 flies were used for each individual experiment, and all crosses were performed at least twice. For cell experiments, all studies were conducted in parallel with vehicle controls in the neighbouring well, for at least 3 wells (biological replicates).

### Western blotting

10 frozen fly head heads or 20 adult retinas were lysed in 50µl ice-cold buffer (20mMTris-Hcl, 100mM Nacl, 1% IGEPAL and 2mM in water) supplemented with Protease Inhibitors (Sigma, MSSAFE-5VL) and Benzonase (Sigma, E1014-24KU) for 25-30 minutes on ice. These samples were centrifuged at 15,000 rpm for 45 minutes at 4 °C. The supernatant from each tube was then mixed with sample buffer (NuPAGE), supplemented with DTT and incubated at 65 °C for 15 minutes. For Oxyblot measurements, supernatants were sequentially treated with 12%SDS, 1X 2-2-dinitrophenyl hydrazine (DNPH) and 1X neutralisation solutions, according to Manufacturer’s instructions (Sigma, S7150), before proceeding to SDS-PAGE analysis.

Proteins were resolved by SDS-PAGE using 4-12%Bis-Tris gels (NuPAGE, Life Technologies) and transferred via electrophoresis to Polyvinylidene difluoride membranes-PVDF (Millipore). Membranes were blocked for 1hour at room temperature in blocking buffer (5%milk in 0.1%triton in PBS/PBST) and incubated overnight at 4 °C in the same buffer containing primary antibodies at 1:1000 dilution. Membranes were washed three times in PBST, and then incubated with secondary antibodies in milk, for 1 hour. Post this step, membranes were again washed three times in PBST and then developed using ECL reagents (Thermo Scientific).

### Cloning of murine and Drosophila ADAM17 constructs and stably expressing cell lines

Murine ADAM17 (mA17), Drosophila ADAM17 (dA17) as well as its mutants each with a PC tag were cloned into the pMOWS vector for retrovirus-based transduction. For the inactive HE mutant of Drosophila ADAM17 (dADAM17) the following single point mutations were introduced: E400H, H409E. HEK293 cells deficient for ADAM17 and ADAM10 (HEK293_A17^-^_A10^-^) (riethmueller) stably expressing the different ADAM17 constructs were generated via the retroviral-based pMOWS/ Phoenix ampho system. 50 µg/ml Zeocin was used as a selection marker.

### Shedding activity assay in HEK cells

Shedding activity of ADAM17 variants was measured by an alkaline phosphatase (AP)–based assay. For this assay 5×10^6^ cells were seeded on a 10-cm dish, and transiently transfected with the ADAM17 substrate IL-1R_II_ fused to an alkaline phosphatase (AP). Transient transfection was performed via the use of Lipofectamine 3000 (Thermo Fischer Scientific). After 24 hours, 2×10^5^ cells/well were transferred into 24-well plates. 24 hour later, cells were treated with only 100 nM PMA, 100 nM PMA combined with 10 µM TAPI-1 or treated with the corresponding volume of vehicle (DMSO). Cells were incubated for 120 minutes at 37 °C. The activity of AP in cell lysates and supernatants was determined by incubating 100 µl AP substrate p-nitrophenyl phosphate (PNPP) (Thermo Scientific) with 100 µl cell lysate or cell supernatant at room temperature followed by the measurement of the absorption at 405 nm. The percentage of AP-conjugated material released from each well was calculated by dividing the signal from the supernatant by the sum of the signal from lysate and supernatant. The data was expressed as mean of at least three independent experiments.

### Statistical analysis of AP assay in HEK cells

Quantitative data are shown as mean with standard deviation (SD) calculated from n = 10 independent experiments. Statistics were conducted using the general mixed model analysis (PROC GLIMMIX, SAS 9.4, SAS Institute Inc., Cary, North Carolina, USA) and assumed to be from a lognormal distribution with the day of experiment conduction as random, to assess differences in the size of treatment effects across the results. Residual analysis and the Shapiro-Wilk test were used as diagnostics. In the case of heteroscedasticity (according to the covtest statement) the degrees of freedom were adjusted by the Kenward-Roger approximation. All p-values were adjusted for multiple comparisons by the false discovery rate (FDR). p<0.05 was considered significant.

### Activity assay in S2R+ cells

Shedding activity of ADAM17 variants was measured by an alkaline phosphatase (AP)–based assay. 24 well plates were first coated with Poly-L-lysine (Sigma). Around 4×10^5^ cells/well were plated onto 24-well plates containing a mix of ADAM17 variants with eiger fused to AP plasmids and transfection reagent (Fugene). Media was changed after 24 hours. 48 hours, post plating cells were treated with only 100 nM PMA, 100 nM PMA combined with 10 µM TAPI-1 or treated with the corresponding volume of vehicle (DMSO). Cells were incubated for 120 minutes at 37 °C. Cell supernatants were collected, the cells were washed in PBS and lysed in 200 µl Trition X-100 lysis buffer. The activity of AP in cell lysates and supernatants was determined by incubating 100 µl AP substrate p-nitrophenyl phosphate (PNPP) (Thermo Scientific) with 100 µl cell lysate or cell supernatant at room temperature followed by the measurement of the absorption at 405 nm. The percentage of AP-conjugated material released from each well was calculated by dividing the signal from the supernatant by the sum of the signal from lysate and supernatant. The data was expressed as mean of at least three independent experiments.

### MDA Analysis of peroxidated lipids

The Lipid Peroxidation (MDA) Assay Kit (Colorimetric/Fluorometric) (Abcam, Cat# ab118970) was used as per the manufacturer’s instructions. 1 million cells or fifteen retinas per sample were dissected in cold 1×PBS, and then transferred to 120 µL MDA lysis buffer with 1 µL BHT. After homogenization, samples were vortexed and centrifuged to remove precipitated protein. 100 µL of the supernatant was added to 300 µL thiobarbituric acid reagent, and incubated at 95 °C for 1 h. 200 µL of standard or sample were added to individual wells of a GREINER 96 F-BOTTOM plate. For fluorometric measurement, signals were collected with a CLARIOStar reader (BMG LABTECH GmbH) (Ex/Em = 532 ± 8/553 ± 8 nm). Three biological replicates were quantified per sample.

### Human induced Pluripotent Stem cells

The iPS cell lines used in this study have all been published previously (SFC840-03-03(Fernandes et al., 2016), SFC841-03-01(Dafinca et al., 2016) SFC856-03-04(Haenseler et al., 2017) and are all available from the European Bank for iPS cells, EBiSC. They were derived in the James Martin Stem Cell Facility, University of Oxford from dermal fibroblasts of healthy donors, who had given signed informed consent for the derivation of iPSC lines from skin biopsies as part of the Oxford Parkinson’s Disease Centre. (Ethics Committee: National Health Service, Health Research Authority, NRES Committee South Central, Berkshire, UK (REC 10/H0505/71). All experiments used cells thawed from large-scale karyotype QCed frozen stocks.

iPS cells were differentiated through a primitive myeloid differentiation pathway to generate primitive macrophage precursors (Buchrieser et al., 2017), followed by maturation according to (Haenseler et al., 2017)to microglial-like cells, seeding at 75,000 cells per well on µ-Plate 96 Well Black-walled imaging plates (Ibidi 89626) precoated for 1 h with geltrex (Life Technologies A1413302), and fed twice-weekly for two weeks with 150 µL microglial medium. Microglial medium was generated to have a final concentration of 1X Advanced DMEM/F12 (Life Technologies, 12634-010), 1X N2 supplement (Life Technologies, 17502-048), 2mM GlutaMAX (Life Technologies, 35050-061), 50µM 2-mercaptoethanol (Life Technologies,31350-010), 50U/ml Pen/Strep (Life Technologies,17502-048), 100ng/ml IL-34 (Peprotech, 200-34), 10 ng/ml GM-CSF(LifeTechnologies, PHC2013).

### Staining and imaging of Human iPS cells

Drug treatments of either 3 μm GW (synthesised by Iris Biotech), 3μm GI (synthesised by Iris Biotech), 10 μm TAPI-1 (Cayman Chemicals) or DMSO (Sigma) were performed by supplementing the media with the above drugs for cells grown on 96 well ibdi Plates, suitable for imaging. Cells were then incubated with either 5nM MitoSOX (Thermo Fischer Scientific) for 30mins at 37^°^C, 200nM C-11 BODIPY (Thermo Fischer Scientific) for 30mins at 37^°^C or 493/503 BODIPY (Thermo Fischer Scientific) and FM 4-64FX (Thermo Fischer Scientific) dyes, for 10mins at 37^0^C. Post-staining cells were almost immediately imaged on the Live Cell Olympus FV1200, maintaining the temperature at 37^°^C throughout the course of the experiment.

### Lipid Droplet Measurements on iPS cells

Lipid droplet numbers and size were measured using an algorithm developed on Image J. Integrated total lipid was calculated as a product of normalised size and total lipid droplet numbers across an individual cell. Normalised LD size was calculated by dividing the size of LDs across all genotypes with that of its corresponding control.

### Image Analysis and compilation

Post-acquisition, all images were analysed on Image J, and subsequently compiled on Adobe Photoshop. The model was compiled on Adobe Illustrator.

## Acknowledgements

We thank Bertrand Mollereau, Viorica L Lastun, Alessia Gelasso and Mike Renne for their insightful comments on the manuscript. We also thank Mike Murphy, Pedro Carvalho and all Freeman lab members for useful discussions; Ni Tang for help with development of the lipid droplet quantifying algorithm; and Pierre Leopold for sharing fly stocks. We particularly thank Dr Errin Johnson and members of the Dunn school EM facility for advice on EM protocols. SM has been funded by HFSP and a non-stipendiary EMBO fellowship. SD has been funded by a Research Fellowship of the German Research Foundation (DU 1582/1-1). This work was primarily supported by a Welllcome Trust Investigator Award (101035/Z/13/Z) to MF.

## Author Contributions

SM and MF conceptualised and wrote the original draft of the paper. SM, CL and MF designed the methodology for experiments. SM and CL conducted all fly related experiments. SM did all the analysis and iPSC cell culture experiments. SD and ID performed cloning of constructs and AP-shedding in HEK cells. SC provided iPS cells and advised on their experimental use.

## Supplemental Information

**Figure S1:**
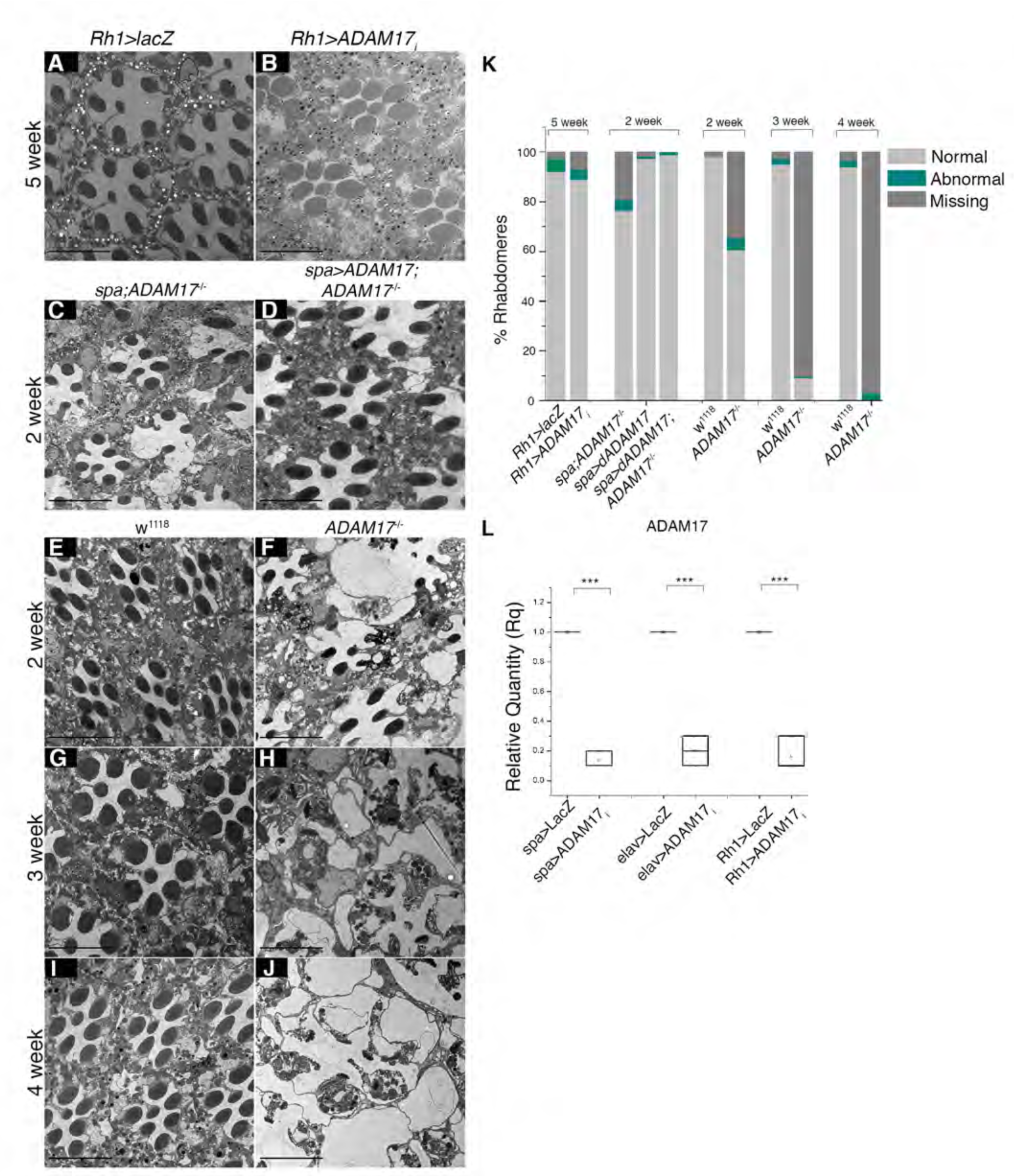
Loss of ADAM17 in PGCs induces age-dependent degeneration: **A-J.** TEM images of adult retinas; (A) over-expressing lacZ in neurons; (B) RNAi for ADAM17 in neurons; (C) *ADAM17*^-/-^ mutant overexpressing WT-ADAM17 in PGCs; (E,G,I) wildtype; or (F,H,J) *ADAM17*^-/-^ mutant retinas at 2 week (E-F), 3 week (G-H) and 4 week (I-J). **K.** Quantitation of the numbers of degenerating photoreceptors from the TEM images corresponding to the genotypes mentioned above; n=180 ommatidia from 3 different fly retinas for each. **L.** qPCR measurements of knockdown efficiency of ADAM17 with *sparkling*, *elav* or *Rh1* GAL4s; n=3 independent biological replicates with 3 technical replicates for each experiment. ***p<.001, **p<.01. Scale bar :10μm

**Figure S2:**
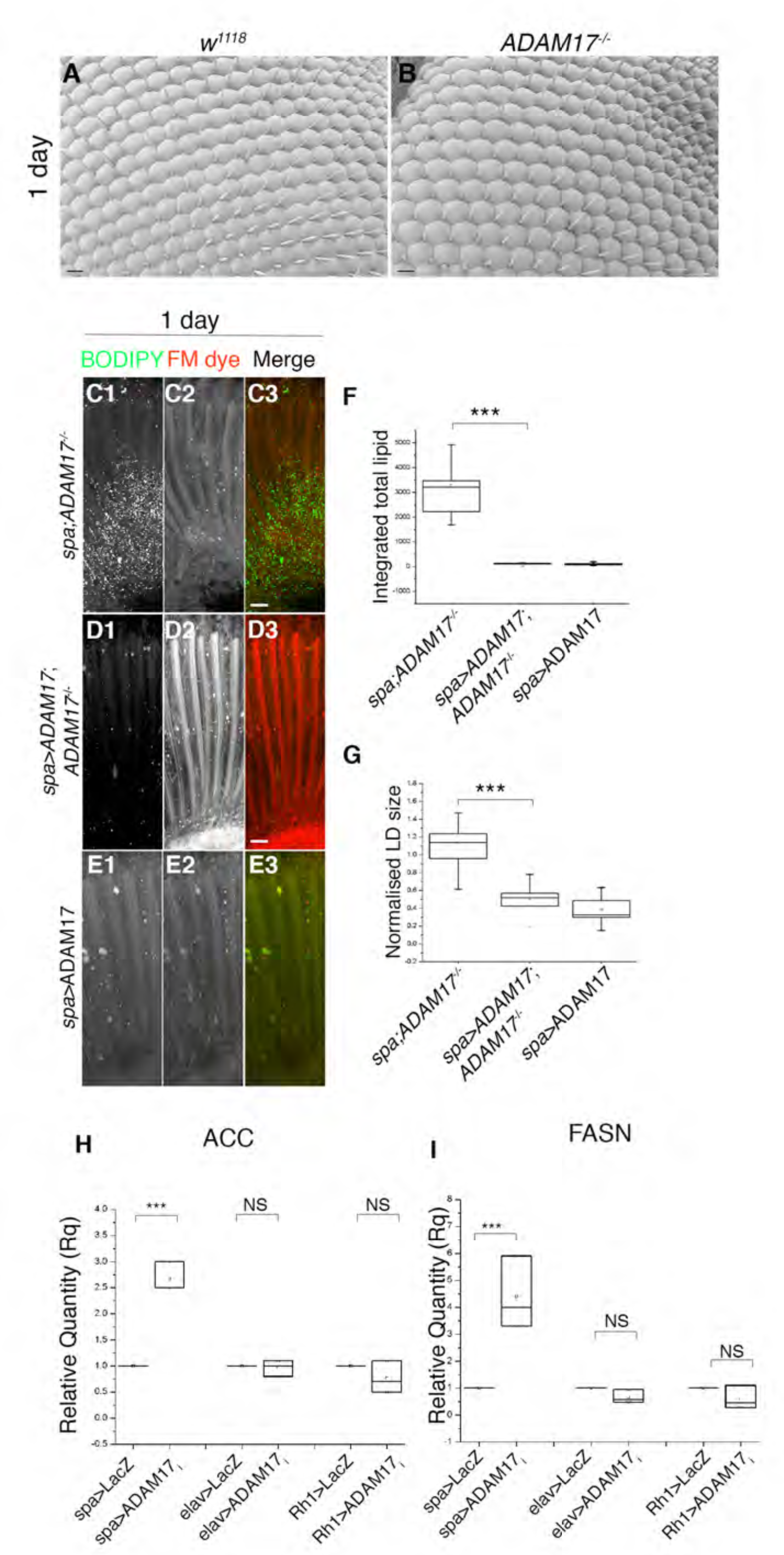
Normal eye development in *ADAM17^-/-^* mutant, rescue of LD phenotype with WT-ADAM17 and expression of lipogenic genes with a knockdown of ADAM17 using different drivers. **A-B.** Scanning Electron Microscopy (SEM) images of the surface of adult retinas corresponding to (A) wild type; and (B) *ADAM17*^-/-^; n=3 flies for each. Scale bar: 10μm **C-E.** Fluorescent images of 1 day old retinas, stained with BODIPY (green) and FM-dye (red); (C) *ADAM17*^-/-^ mutant; (D) glial specific overexpression of ADAM17 in *ADAM17*^-/-^ mutant; or (E) glial specific overexpression of ADAM17 in a wild-type background. **F-G.** Quantitation of BODIPY staining (shown in C-E) depicted as (F) integrated total lipid and (G) normalised lipid droplet size; n=10 for each genotype. Scale bar:10 μm. **H-I.** qPCR measurements of lipogenic transcripts of ACC and FASN across different knockdowns of ADAM17 with *sparkling, elav or Rh1* GAL4s; independent biological replicates with 3 technical replicates for each experiment. ***p<.001

**Figure S3:**
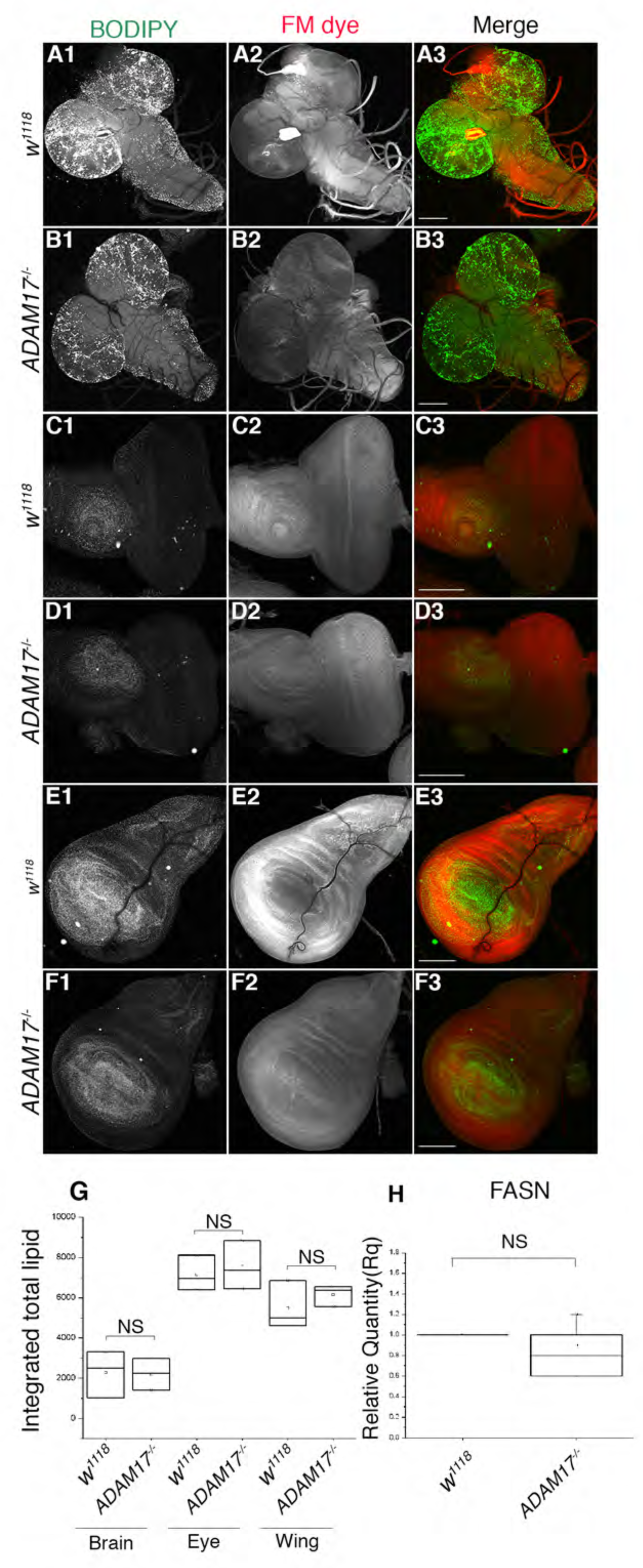
Loss of ADAM17 does not affect LD in larval tissues. **A-F.** Fluorescent images of either wild type or *ADAM17*^-/-^ mutant larval (A-B) brain; (C-D) eye imaginal disc; and (E-F) wing imaginal disc, stained with BODIPY (green) and FM-dye (red). **G.** Quantitation of BODIPY staining (shown in A-F) depicted as integrated total lipid; n=10 for each genotype. **H.** FASN mRNA transcript levels measured by qPCR in wild-type and *ADAM17*^-/-^ larvae; n=4 for each.

**Figure S4:**
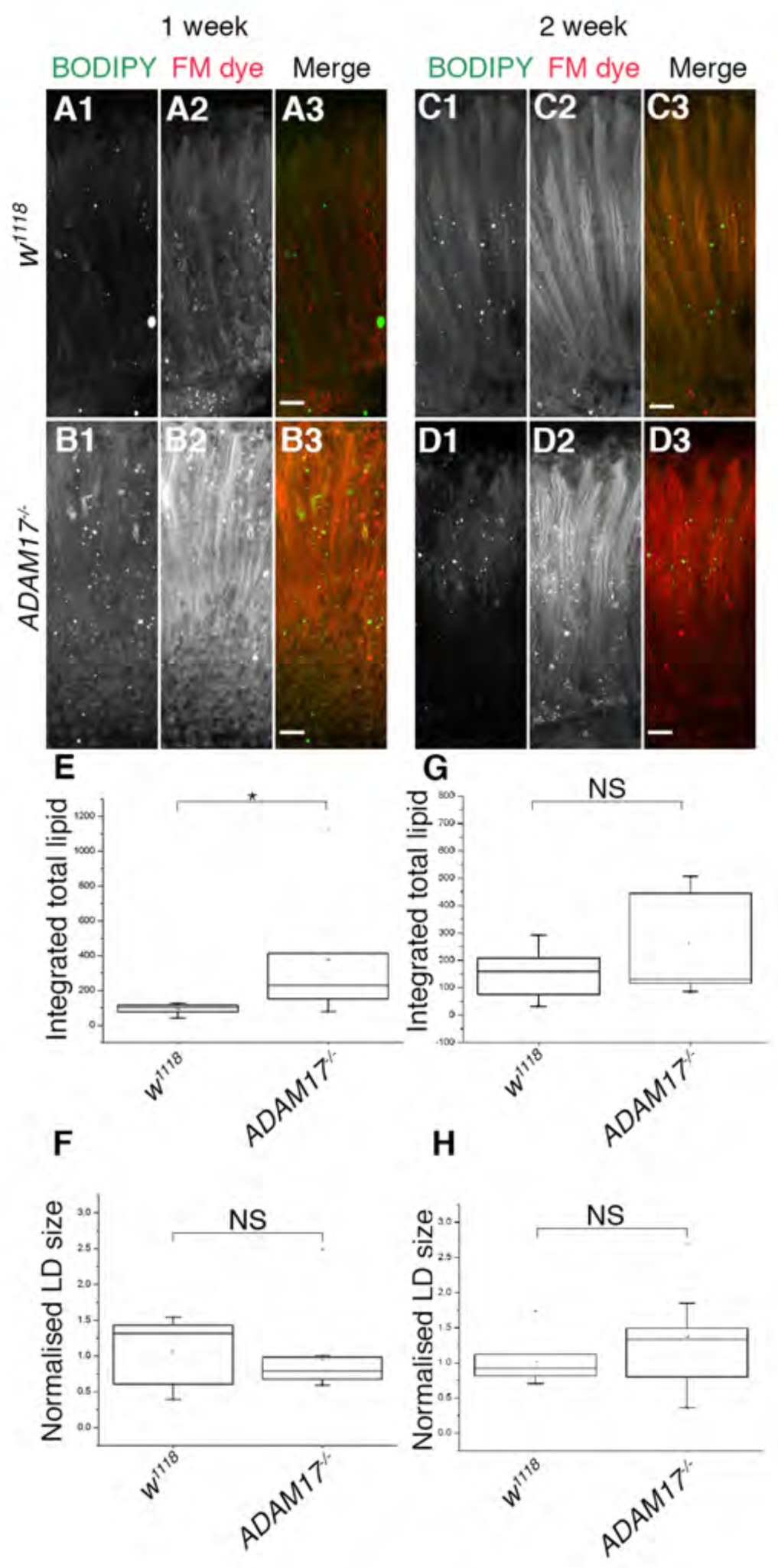
Age-dependent clearance of LD in ADAM17 mutant retinas. **A-D.** Fluorescent images of 1week old (A) wild type and (B) *ADAM17*^-/-^; 2 week old (C) wild type and (D) *ADAM17*^-/-^ mutant retinas labelled with BODIPY (green) and FM dye (red). **E-H.** Quantitation of BODIPY staining (shown in A-D), depicted as integrated total lipid (E, G) and normalised lipid droplet size (F, H); n=10 for each genotype. *p<.05

**Figure S5:**
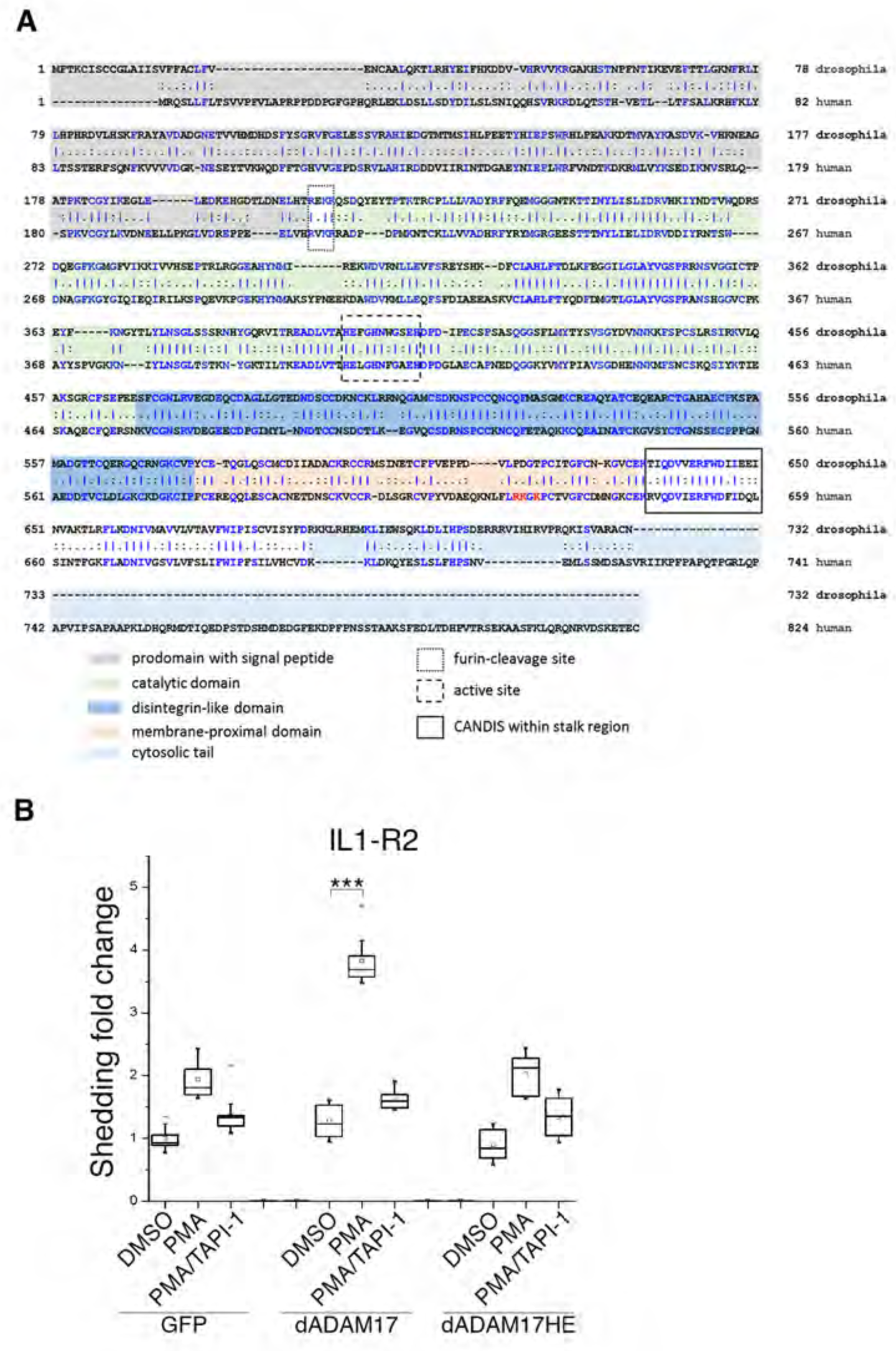
Sequence comparison and AP-shedding assays of Drosophila ADAM17. **A.** Sequence alignment of Drosophila ADAM17 with its human counterpart. **B.** AP-shedding assays performed for IL1-R2 in HEK cells, lacking both human ADAM17 and ADAM10, with either GFP control, full length Drosophila ADAM17, or an activity dead mutant of Drosophila ADAM17, in the presence of either PMA or PMA and TAPI-1 (DMSO is used as control); n=10 for each condition. ***p<.001

**Figure S6:**
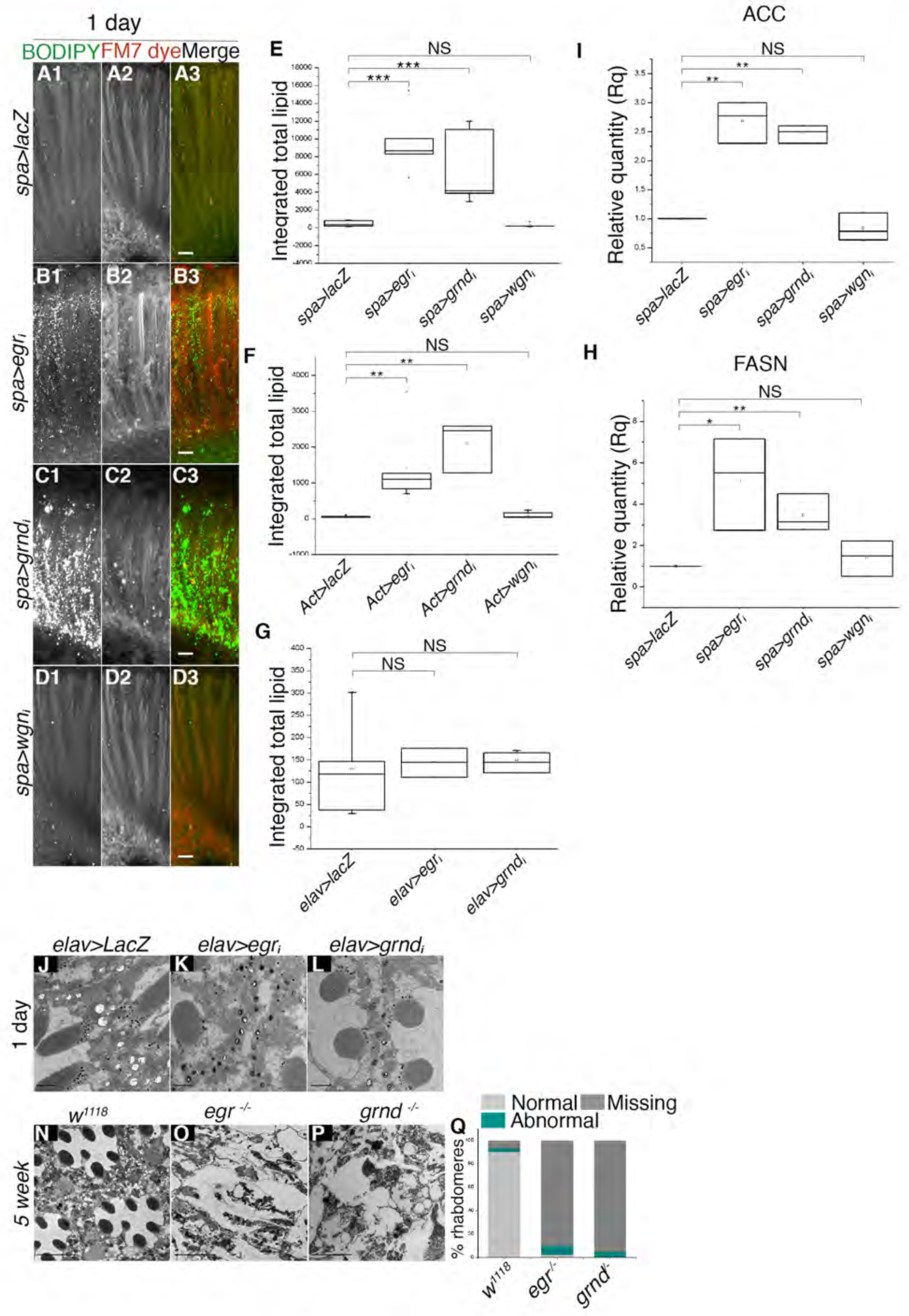
Loss of eiger and grindelwald, but not wengen specifically in PGC leads to abnormal LD accumulation and age-associated retinal degeneration. **A-D.** Fluorescent images of 1 week old retinas labelled with BODIPY (green) and FM dye (red); overexpressing (A) *lacZ*; (B) *eiger* RNAi; (C) *grindelwald* RNAi or (D) *wengen* RNAi specifically in PGC; n=10 for each genotype. Scale bar: 10 μm. **E.** Quantitation of BODIPY staining (observed in A-D) shown as integrated total lipid count. **F-G.** Quantitation of BODIPY staining in retinas overexpressing *lacZ*, *eiger* RNAi, *grindelwald* RNAi or *wengen* RNAi either throughout the retina (F); and the neurons (G); n=10 for each (Fluorescent images not shown). **H-I.** mRNA transcript levels of ACC and FASN measured by qPCR in retinas expressing *lacZ*, *eiger* RNAi, *grindelwald* RNAi or *wengen* RNAi in PGC; n=3 independent biological replicates with 3 technical replicates for each experiment. **J-L.** TEM images of 1 day old retinas expressing (J) *lacZ*; (K) *eiger* RNAi; (L) and *grindelwald* RNAi specifically in neurons. **N-P.** TEM images of 5 week old retinas of (N) wild type; (O) *eiger;* and (P) *grindelwald* mutants. **Q.** Quantitation of the numbers of degenerating photoreceptors observed in the TEM images corresponding to the genotypes mentioned above in N-P; n=180 ommatidia from 3 different fly retinas for each. Scale bar :10μm. ***p<.001, **p<.01, *p<.05

**Figure S7:**
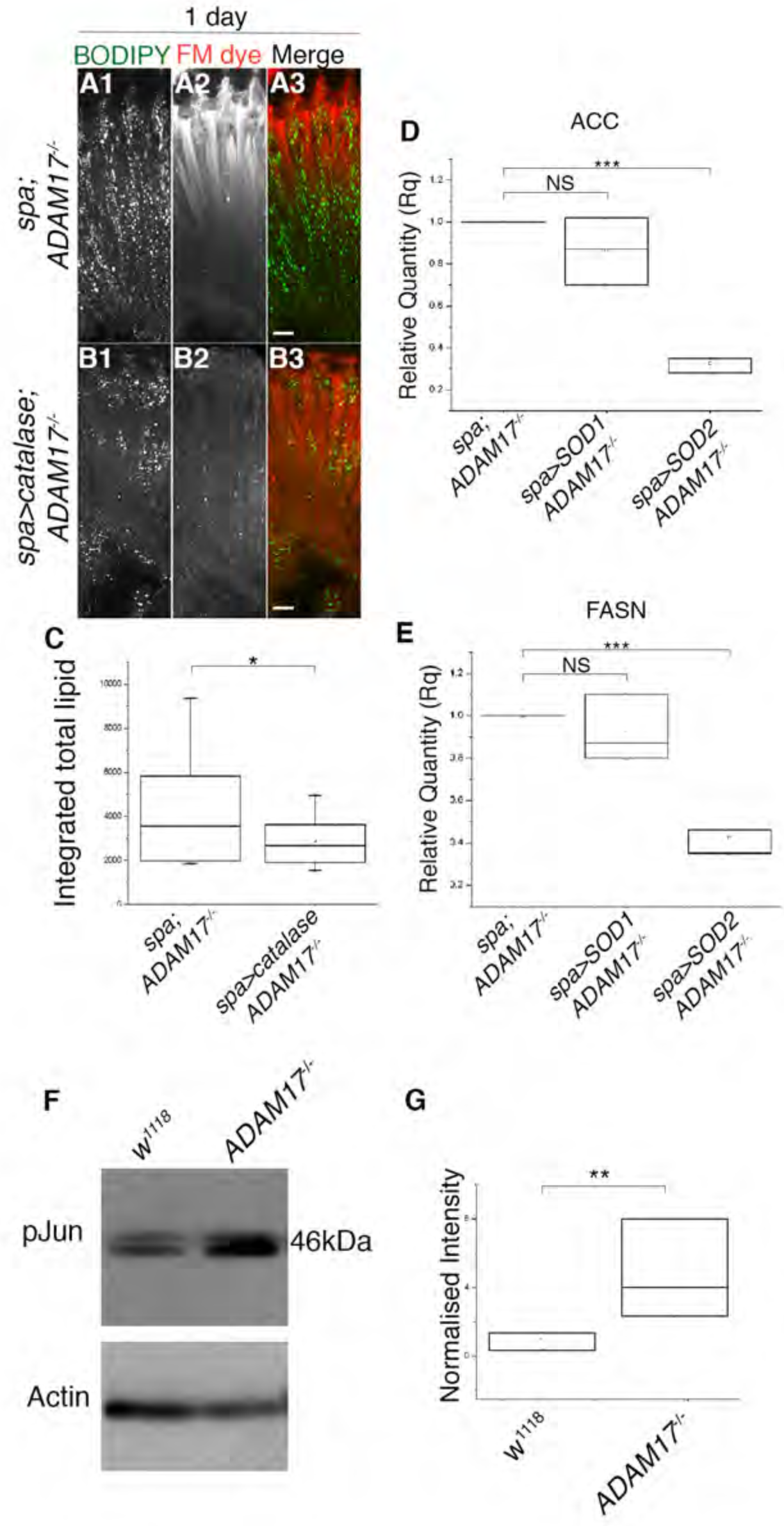
Lipid droplet counts and lipogenic transcript levels in glial specific overexpression of catalase, SOD1 or SOD2. **A-B.** Fluorescent images of retinas stained with BODIPY (green) and FM dye (red) corresponding to (A) *ADAM17*^-/-^; and (B) glial specific overexpression of catalase in *ADAM17*^-/-^ background; n=10 for each genotype. Scale bar: 10μm. **C**: Quantitation of BODIPY staining (shown in A and B), depicted as integrated total lipid. **D-E.** qPCR measurement of mRNA transcript levels of ACC and FASN from *ADAM17*^-/-^ retinas and retinas overexpressing either SOD1 or SOD2 within glial cells in an *ADAM17*^-/-^ background. **F.** Western blot analysis of phosphorylated Jun (pJun) and actin levels from head lysates of wild type and *ADAM17*^-/-^mutants. **G**. Averaged intensity levels of pJun normalised to actin, as inferred from western blots. n=5 different biological replicates. ***p<.001, *p<.05

**Figure S8:**
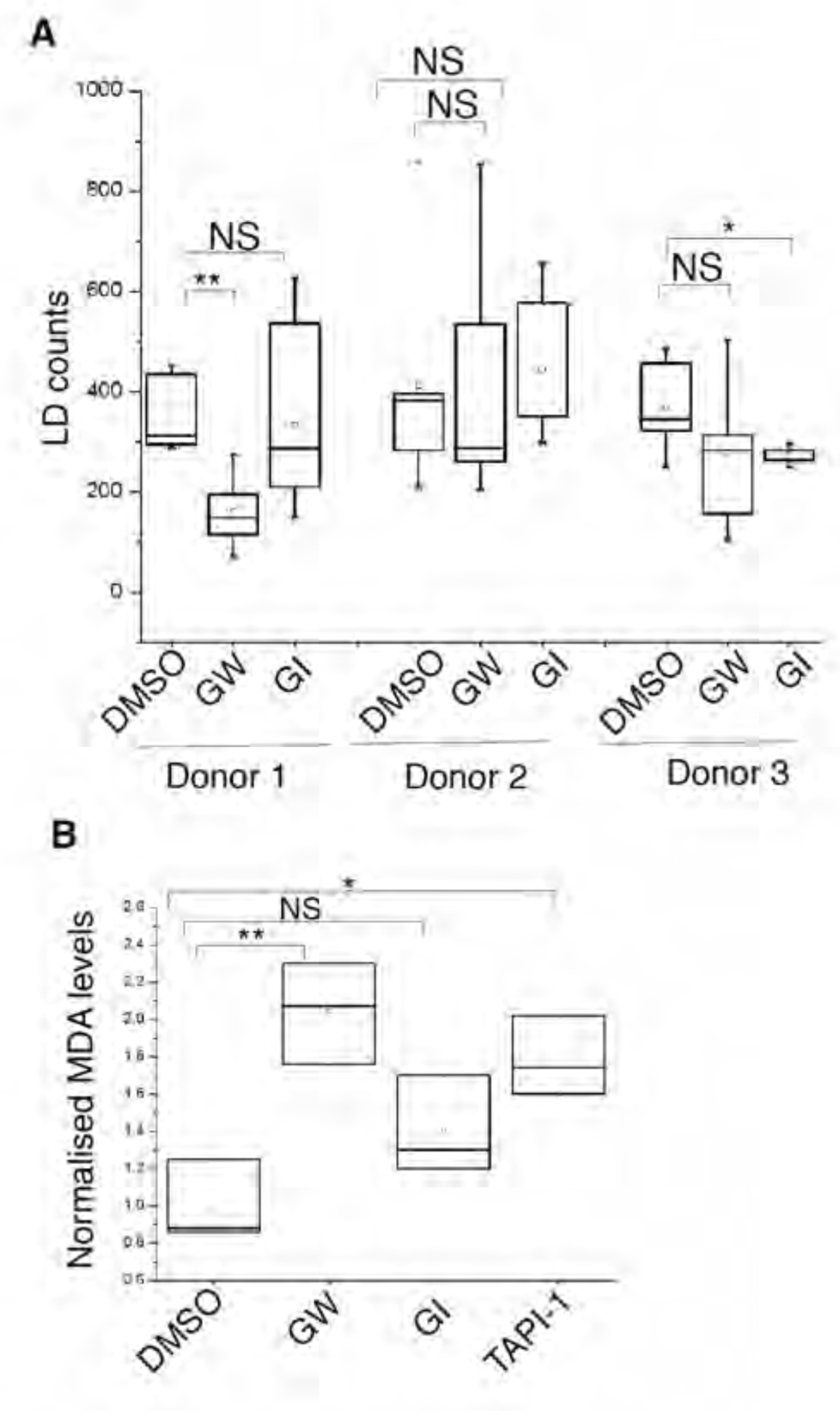
LD counts and MDA amounts for iPSC microglia cells treated with inhibitors for ADAM17 and/or ADAM10. **A.** LD counts for iPSC derived microglia-like cells from 3 different donors treated with either DMSO, GW or GI for a period of 24hours; n=10 cells for each treatment, per donor. **B.** Levels of Malondialdehyde (normalised to DMSO controls) in cell lysates from Donor No.2 treated with either DMSO, GW, GI or TAPI-1 for 24 hours; n=3 biological replicates., **p<.01, *p<.05

### Fly genotypes

#### Fig. S1

*;+/+*;*Rh1-GAL4*/*UAS-lacZ/+*

*;+/+*;*Rh1-GAL4*/ *UAS-ADAM17_i_/+spa-GAL4*;+/+;*ADAM17^-/-^*

*spa-GAL4*;UAS-ADAM17-WT/+;*ADAM17^-/-^*

*spa-GAL4*;UAS-ADAM17-WT/+;

*w^1118^*

*;;ADAM17^-/-^*

#### Fig. S2

*w^1118^*

*;+/+;ADAM17^-/-^*

*spa-GAL4*;+/+;*ADAM17^-/-^*

*spa-GAL4*;UAS-ADAM17-WT/+;

*spa-GAL4*;UAS-ADAM17-WT/+;*ADAM17^-/-^*

#### Fig. S3

*w^1118^*

*;+/+;ADAM17^-/-^*

#### Fig. S4

*w^1118^*

*;+/+;ADAM17^-/-^*

#### Fig. S6

*spa-GAL4*;+/+;*UAS-LacZ/+*

*spa-GAL4*;+/+;*UAS-eiger_i_/+*

*spa-GAL4*;+/+;*UAS-grnd_i_/+*

*spa-GAL4*;+/+;*UAS-wgn_i_/+*

;+/+;*Actin-GAL4/UAS-LacZ*

*+/+;*;*Actin-GAL4*/*UAS-eiger_i_*

*;+/+*;*Actin-GAL4*/*UAS-grdn_i_*

*;+/+*;*Actin-GAL4*/*UAS-wgn_i_*

*;elav-GAL4*/+;*UAS-LacZ/+*

*;elav-GAL4*/+;*UAS-egr_i_/+*

*;elav-GAL4*/+;*UAS-grnd_i_/+*

*w^1118^*

*egr^-/-^*

*grnd^-/-^*

#### Fig. S7

*w^1118^*

*spa-GAL4*;;*ADAM17^-/-^*

*spa-GAL4*;UAS-SOD2/+;*ADAM17^-/-^*

*spa-GAL4*;UAS-SOD1/+;*ADAM17^-/-^*

*spa-GAL4*;UAS-catalase/+;*ADAM17^-/-^*

## Notes

#### Summary of Updates

References corrected

